# The genome of the vining fern *Lygodium microphyllum* highlights genomic and functional differences between life phases of an invasive plant

**DOI:** 10.1101/2025.03.06.640867

**Authors:** Jessie A. Pelosi, Ruth Davenport, Li-Yaung Kuo, Levi N. Gray, Anthony J. Dant, Emily H. Kim, Fay-Wei Li, Katrina M. Dlugosch, Trevor J. Krabbenhoft, W. Brad Barbazuk, Emily B. Sessa

## Abstract

Functional and genomic studies on the differences between gametophyte and sporophyte life phases remain scarce, yet unraveling these dynamics is crucial to understanding the biology of plants and the success of each phase under different environments. Here, we provide a novel reference genome for the highly invasive fern *Lygodium microphyllum* and compare the transcriptomic and epigenomic landscapes of the gametophyte and sporophyte life phases. We found differential regulation of developmental genes (homeobox and MADS-box clades) and usage of alternative isoforms that may play a role in the genomic determination of the haploid and diploid life stages. We further generated the first base pair-resolution methylome of a fern gametophyte, and determined that methylation patterns are remarkably similar between vegetative tissues despite their morphological and functional differences. By examining the physiological and transcriptomic responses of gametophytes and sporophytes to freezing stress, the most likely abiotic factor preventing further expansion of this invasive species, we show that life phases and tissues use alternative molecular pathways to respond to this stressor, underscoring the need to incorporate both life phases when developing effective mitigation strategies. These new genomic resources fill a gap in our understanding of fundamental plant biology and inform invasive species research.

**SIGNIFICANCE:** All land plants undergo an alternation of generations between haploid gametophyte and diploid sporophyte life phases. How these disparate life phases are generated from a single genome, and the functional implications of these differences for plant success, is largely unknown. Moreover, understanding life-phase-specific differences in rapidly evolving populations, such as invasive species, is critical to developing effective management strategies. We assembled a chromosome-level genome and generated epigenomic and transcriptomic resources from gametophyte and sporophyte phases of the invasive fern *Lygodium microphyllum,* which is estimated to cost the U.S. more than $2 million annually. We highlight transcriptomic, epigenomic, and physiological variation between life phases and find support for the crucial role of the often overlooked gametophyte in the invasion process.

## MAIN

All land plants alternate between multicellular diploid sporophyte and haploid gametophyte life phases (typically referred to as the “alternation of generations” life cycle). Variations in this life cycle mark pivotal transitions in the evolution of plants (e.g., the transition from gametophyte-dominance in bryophytes to sporophyte-dominance in vascular plants), but very little is known about how a single underlying genome generates these radically disparate stages. Ferns are an excellent system for unraveling these processes as they have free-living, nutritionally-independent gametophytes and sporophytes (1). Previously, researchers have found minimal but distinct differences in gene expression level between fern life stages (e.g., up to 90% of genes had no significant differences in expression level (2–4)), but the regulation and functional roles of these genes remains a crucial gap in our knowledge. This is in part because genomic resources for this clade still lag behind other major lineages despite recent publications of high-quality fern genomes (e.g., (3, 5–8).

The gametophyte has long been thought of as a fragile handicap in the fern life cycle (9, 10), and has largely been overlooked by researchers. The gametophyte generation, however, is the site of sexual reproduction and is thus pivotal for the establishment of sporophyte populations and will shape evolutionary trajectories (11, 12). Furthermore, physiological evidence suggests that the haploid life phase of ferns may in fact be more hardy than the sporophyte, able to withstand extreme desiccation (13, 14), freezing (15, 16), drought (17–19), and heat (19). These differences highlight a largely neglected fundamental aspect of organismal plant biology, with profound functional implications for our understanding of the processes that shape the niche, distribution, and evolution of ferns (reviewed by (20)).

The application of high-throughput sequencing to explore the genomic underpinnings of these functional differences is under-explored yet has provided valuable insights in other groups, including improving our understanding of the regulation of the alternation of generations in algae (21), bryophytes (22, 23), and angiosperms (24), and revealing large-scale genomic re-organization associated with transitions in the life cycles of *Arabidopsis* (*25*) and some animals (26). Of particular interest is how differences between life stages play out in rapidly-evolving systems such as invasive species, which pose an unprecedented threat to ecological, environmental, and economic systems globally (27). Recent estimates place the economic costs of biological invasions at over $100 billion annually world-wide (28) and over $20 billion annually in the United States alone (29). Given the functional variation between life stages in many organisms, determining the relative roles of the gametophyte and sporophyte generations in the invasion process of ferns may have implications for invasion biology generally, including in the development of effective management strategies.

Here, we present a high-quality, chromosome-scale reference genome for the invasive fern *Lygodium microphyllum* (Fig. 1). This species is a highly successful vine with indeterminate rachis growth (30) and extensive rhizomatous proliferation, which facilitates the formation of dense mats that smother and shade native ecosystems, disrupt natural fire regimes (31, 32), and alter native community composition through canopy collapse (33). In its invasion of the United States, *Lygodium microphyllum* is currently restricted to southern Florida, as freezing is widely recognized as range-limiting for its invasive populations (34, 35). We use this new genomic resource to explore differences in life stages at multi-omic scales from epigenomes to transcriptomes, and highlight how different physiological and genetic responses of the gametophyte and sporophyte to cooling and freezing may have important implications for management in a changing climate.

**Figure 1.**
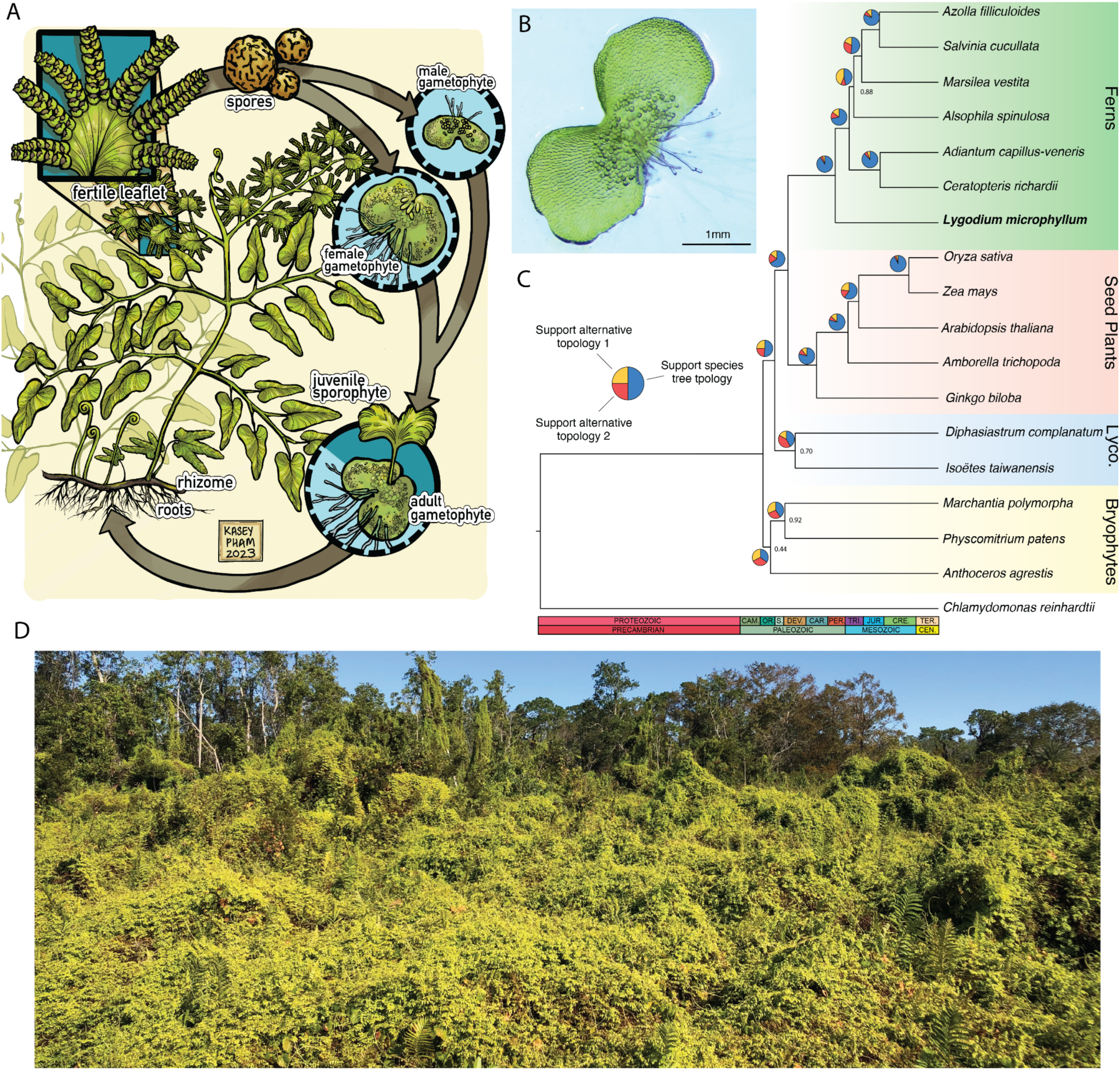
Genome assembly of *Lygodium microphyllum.* **A)** The life cycle and unique morphology of *Lygodium microphyllum* (illustration by Kasey Pham). The sporophyte-dominated life cycle alternates between the diploid sporophyte and haploid gametophyte phases. The vining habit is facilitated by dormant pinnae buds and circumnutation, depicted here at the apex of the leaf. **B)** Antheridate gametophyte of *L. microphyllum*; scale bar is 1 mm.. **C)** Phylogenetic placement of *L. microphyllum* within the Viridiplantae on a time-calibrated tree. Major lineages of land plants are delineated by background color. Pie charts at nodes show the proportion of gene trees that support each topology (species tree topology or the two alternative topologies). **D)** Field site heavily impacted by *L. microphyllum* invasion in Florida (the light green plant dominating the fore and midground).

## RESULTS AND DISCUSSION

### Genome Assembly and Annotation

The size of the *Lygodium microphyllum* genome has been estimated to be between 4.2-5.1 Gb, with low heterozygosity (0.106%) based on k-mer approaches (Fig. S1), which are similar to flow cytometry-based estimates of 5.56 Gb (36). Using a combination of corrected Oxford Nanopore (37x coverage) and Illumina (60x coverage) reads, we generated a contig-level assembly that was 4.75 Gb in total length with a contig N50 of 17.37Mb (Table 1, S1). By integrating chromatin conformation capture data (Dovetail Omni-C; 73x coverage), we scaffolded the genome to the chromosome-scale (Fig. S2), with a total of 4.61 Gb (96.94%) anchored on 30 pseudomolecules representing the 30 chromosomes in the *L. microphyllum* karyotype (37, 38). The final assembly contained 952 scaffolds with a scaffold N50 of 155.01Mb (Table 1), and a plastome assembly of 163kb (Fig. S3). Of the 30 chromosome-level scaffolds, we identified high proportions of telomeric repeats (AAACCCT) on at least one end for 28 scaffolds, and on both ends of 11 scaffolds (Fig. S4). The genome assembly was highly complete, with 98.11% single and duplicated intact BUSCO genes (Viridiplantae ODB10, *n*=425, Table 1). The mapping rate of genomic short-read Illumina data was over 99%, indicating a highly complete and accurate assembly.

**Table 1.** Genome assembly and annotation statistics.

| **Metric** | **Value** |
| --- | --- |
| Final Assembly Size | 4.75Gb |
| Number of Contigs | 1,801 |
| Contig N50 | 17.35Mb |
| Number of Scaffolds/Unplaced Contigs | 952 |
| Number of Chromosome-Level Scaffolds | 30 |
| Scaffold N50 | 155.01Mb |
| Repetitive Content | 78.59% |
| Genome BUSCO (Viridiplantae ODB10, *n*=425)<br>[No. Single Complete, No. Duplicated Complete] | 98.11% [409S, 8D] |
| Number of Predicted Protein-Coding Genes | 22,827 |
| Proteome BUSCO (Viridiplantae ODB10, *n*=425)<br>[No. Single Complete, No. Duplicated Complete] | 98.80% [400S, 20D] |
| Proteome OMArk (Tracheophyta, *n*=6460) | 91.97% |

As with most plants, the majority of the *Lygodium* genome was composed of repetitive elements: 78.59% of the genome was occupied by repeats. Most of these elements were classified as long terminal repeats (59.97% of the genome) in the Gypsy (33.77%) and Copia (17.94%) families. A small proportion of the repeats were DNA transposons (3.91% of the genome), largely classified as CMC-EnSpm elements (3.30%; see Table S2). On average, repeat density was high with approximately 1,092 repetitive elements/Mb of sequence, while gene density was low with 4.89 genes/Mb. The structural gene annotation using nine tissue types (Table S3) identified 22,827 genes, which is slightly less than the number of genes annotated in other diploid ferns genomes (3, 5, 7). Nearly all genes (22,625, 99.11%) were located on the thirty chromosome-level scaffolds. We found that the proteome was highly complete, with 98.80% BUSCO and 91.27% OMArk scores (Table 1). Gene metrics (e.g., length, number of exons) are similar to those in other published fern genomes (Tables S4-7).

Several recent studies of transcriptomic datasets have identified a putative whole genome duplication (WGD) using *K_S_* -based and gene tree duplication approaches shared by nearly all leptosporangiate ferns, likely dating to sometime in the Permian or Triassic (39–43). We do not find evidence of this ancient polyploidy event in the *Lygodium* genome, with a clear lack of conserved synteny and no burst of duplicates in the *K_S_* plot (Fig. 2). This may not be surprising given that angiosperm genomes undergo rapid rearrangement and loss of duplicate genes that would mask ancient WGDs (44). How rediploidization occurs in ferns, however, is largely unknown and different lineages support a rediploidization continuum from remarkably well-conserved synteny in *Alsophila spinulosa* (*6*) to massive genomic reorganization in *Ceratopteris richardii* (7) and *Adiantum capillus-veneris* (*3*). It appears that there is a fundamental biological difference in this process between flowering plants and ferns, with some hypotheses suggesting that ferns primarily silence or pseudogenize duplicated genes but retain chromosomal material (45–48). Further study on genic fractionation and chromosomal retention will be required to determine why ferns differ so dramatically from other plant lineages in the rediploidization process.

**Figure 2.**
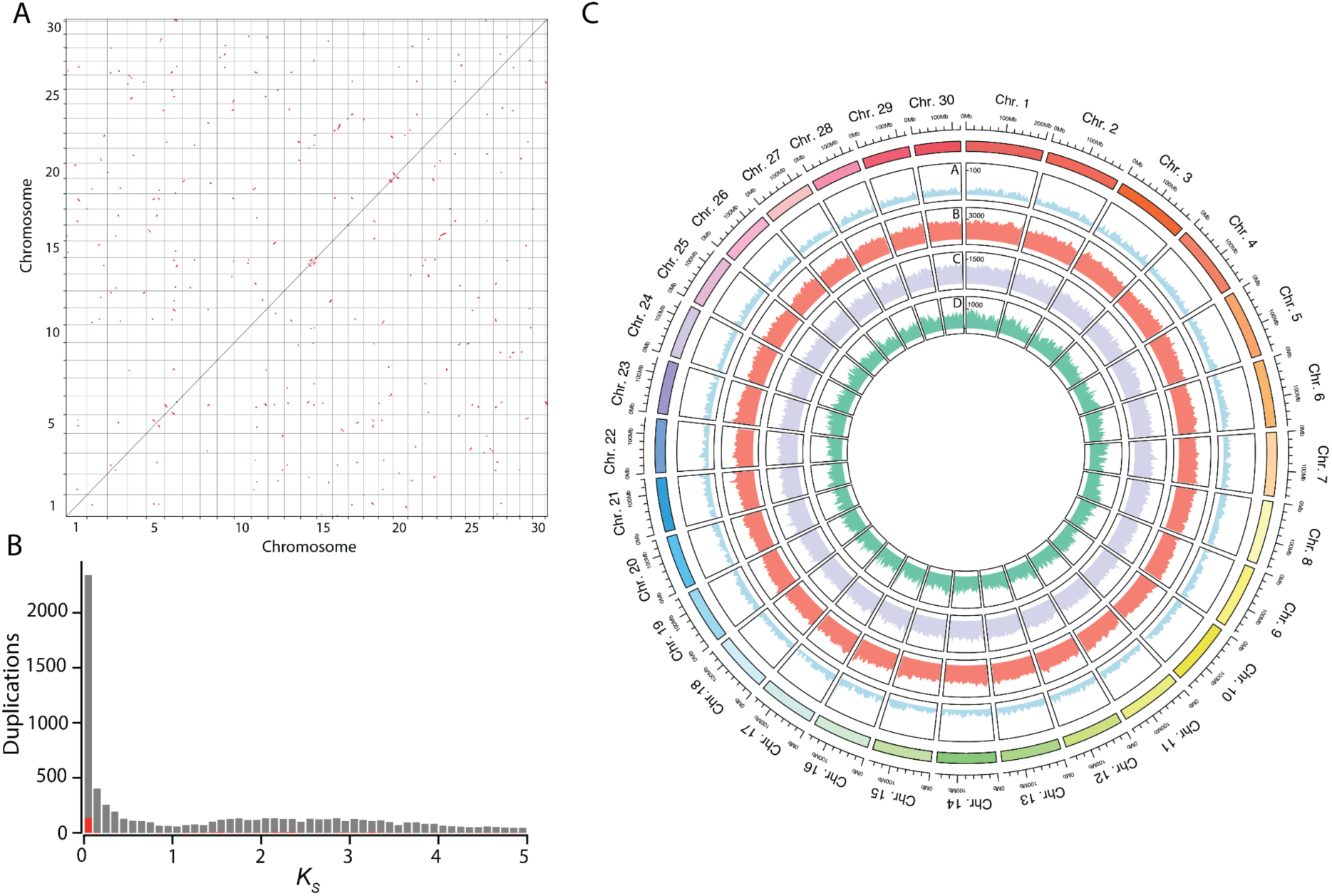
No evidence of whole genome duplication from the *Lygodium microphyllum* genome. **A)** Dot plot of self-self synteny. **B)** Plot of synonymous substitution rates (*K_S_*) for all duplicated genes (gray) and syntenic duplicates (red). Analyses of synteny (A) and *K_S_* (B) do not support a recent whole genome duplication in *L. microphyllum*. **C)** Circos plot of the genome assembly of *L. microphyllum,* showing attributes of the 30 chromosome-level scaffolds. From the outside in, tracks are A) gene density, B) total repeat density, C) Gypsy family repeat density, and D) Copia family repeat density . Densities were calculated using a 5Mb window size.

### Transcriptomic Landscapes of Fern Life Phases

How a single genome can produce two vastly different life phases has long fascinated, and continues to challenge, plant biologists (49). The differential regulation of the plant developmental program is likely a central player, although the mechanism through which this happens has yet to be determined. To compare the transcriptomic landscape of the gametophyte and sporophyte life phases, we mapped RNAseq reads generated from sterile leaves (hereafter ‘sporophyte’ in this section) and pooled immature and mature gametophytes grown under the same ambient conditions to the reference genome and compared their expression levels and splicing patterns.

A total of 1893 (9.5%) genes had significantly different expression levels between life phases: 5.7% of all genes (1136) were upregulated and 3.8% (757) were downregulated in the gametophyte relative to the sporophyte. Our findings are similar to those in a transcriptome-based study of gene expression (9.8% of genes had significantly different expression levels between gametophyte and sporophytes in *Polypodium amorphum* (2)), and genome-based studies (12.6% and 5.8% of genes were differentially expressed in *Adiantum capillus-veneris* (3) and *Ceratopteris richardii* (7), respectively). Most pathways that were over-represented in differentially expressed genes were related to metabolism (Table S8, S9). Of particular interest is the diterpenoid biosynthesis pathway that was enriched in upregulated genes in the gametophyte (8.61 fold enrichment [F.E.], adjusted *P*=0.0019; Fig. S5), which is involved in the generation of gibberellins. In particular, reactions in the gibberellin A12 and A4/1 biosynthesis modules were enriched within this pathway. Gibberellins are regulators of plant growth and development (50), and have been implicated in antheridiogen production in the fern gametophyte (51, 52). The expression of genes such as mono- and dioxygenases (e.g., *KAO2, GA3, GA1*), which produce enzymes that catalyze the reactions from precursor molecules to several GAs, was greater in the gametophyte than in the sporophyte. Both GA_9_ and GA_4_ are vital for sex determination through the antheridiogen pathway detailed in *L. japonicum* (51) and upregulation of genes is expected as the gametophyte matures. Our samples contained a mixture of immature and mature *Lygodium* gametophytes, but were a similar size to those that upregulated *KAO* and *GA20OX* in *L. japonicum* (*51*). Downregulated genes in the gametophyte were enriched for the flavonoid biosynthesis pathway (10.53 F.E., adjusted *P* = 1.77 x10^-3^; Fig. S6), including chalcone synthase (*CHS*) and flavonoid 3’-monooxygenase (*TT7*) genes.

Secondary compounds, such as flavonoids, play an important role in defenses against herbivory in ferns (53). While chemical responses in the sporophyte have shed light on fern resistance to herbivory (3), the gametophyte has largely been ignored in this regard. Further investigation is needed to determine whether the lower expression levels of flavonoids indicate as lower constitutive expression of herbivore resistance compounds in the gametophyte, especially as there are active biocontrol programs implemented to mitigate invasive populations of *L. microphyllum* (54–57) that could take advantage of herbivory on the gametophyte life stage. Alternatively, flavonoids protect plants from UV radiation (58), and a study of *CHS* mutants in *Arabidopsis* that lack flavanols identified that the loss of these compounds resulted in greater susceptibility to UV (59). (60) identified several upregulated flavonoid biosynthesis genes in *Adiantum* gametophytes as a response to high light levels. The higher expression in the *Lygodium* sporophyte at ambient light may suggest that leaves make heavy use of this strategy for protection from UV in this species, while the gametophyte reduces this investment.

We further examined two classes of genes known to be involved in directing plant development. Using a phylogenomic approach, we identified 46 homeobox genes in the *Lygodium* genome annotation (Figs. 3, S7, Table S10). Interestingly, we identified a fern-specific clade of putative homeobox genes that fell sister to the major clade of HDZIP-I genes, similar to the findings in (61). Six homeobox genes had significantly different expression levels in the gametophyte and sporophyte. All had higher expression in the sporophyte than the gametophyte, except for a single gene (g3472, homolog of *PROTODERMAL FACTOR 2 (PDF2)*). In *Arabidopsis, PDF2* is essential for epidermal cell differentiation (62) and downstream stomatal development (63), although stomata are absent in fern gametophytes (20) suggesting that this gene may play another role in gametophyte development. In *Lygodium,* the downregulated genes in the gametophyte are members of the KNOX-II, BEL, and fern-specific HDZIP families, which appears to be a consistent pattern across millions of years of fern evolution, with significantly lower expression of KNOX-II genes in the gametophytes of *Adiantum capillus-veneris* and *Ceratopteris richardii* (Fig. 3D). KNOX genes are involved in sporophyte meristem development and leaf architecture in *Ceratopteris* (64), although few developmental studies are available for ferns. While we cannot speculate on the functions of the fern-specific HDZIP genes, the lower expression levels of BEL and KNOX-II genes aligns with previous findings in bryophytes (e.g., *Anthoceros* (*22*)). Specifically, KNOX-II genes in the sporophyte are required to maintain sporophyte cell fates in *P. patens* (*65*) and *M. polymorpha* (*66*). Our findings highlight a conservation of expression levels in these homeobox genes across >250 my of fern evolution, and perhaps >500 my of land plant evolution.

**Figure 3.**
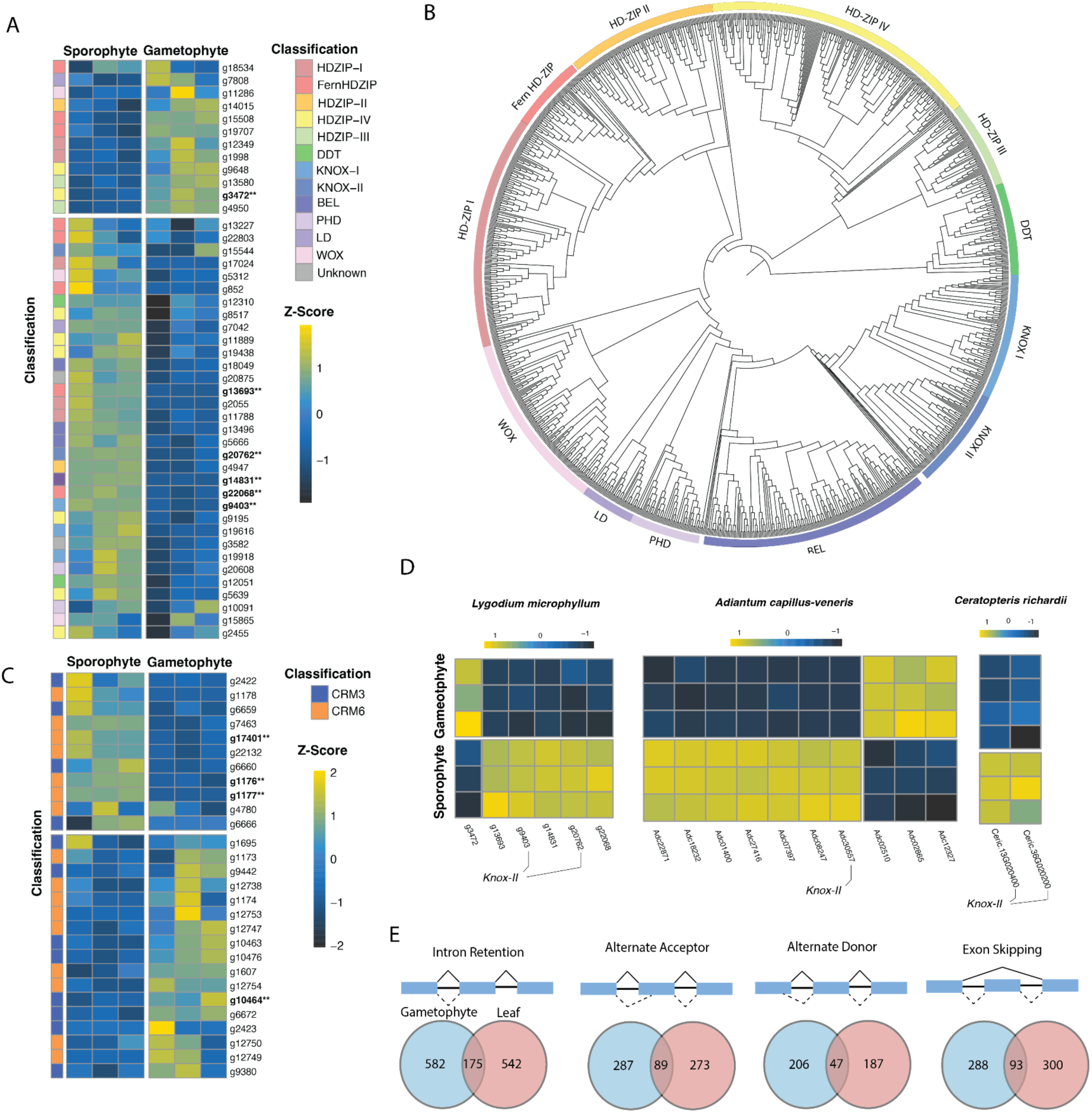
Transcriptomic landscape of gametophyte and sporophyte life phases. **A)** Heatmap of Z-scores of expression levels of homeobox genes identified in the *Lygodium microphyllum* genome. Significantly differentially expressed genes (log2FoldChange > 2, *P* < 0.05) are bolded and denoted with two asterisks (**). **B)** Phylogeny of homeobox genes depicting the relationships of families across land plants. Classification colors are as in **(A)**. **C)** Heatmap of Z-scores of expression levels of CRM-3 and CRM-6 MADS-box genes identified in *Lygodium*. Significantly differentially expressed genes denoted with two asterisks (**) as in **(A)**. **D)** Significantly differentially expressed genes between sterile leaf and gametophyte tissue in three homosporous ferns. KNOX-II consistently has lower expression in the gametophytes of all three species. **E)** The four classifications of alternative splicing events considered in this study, with numbers indicating unique events in the gametophyte (blue) and leaf (red) tissues, and that are shared (intersections).

We also identified a total of 50 MADS-box genes in *Lygodium* (Fig. S8, Table S10). Of these, seven were significantly differentially expressed between sporophyte and gametophyte tissue; two genes (one in the CRM3 clade and one in MIKC* clade) were upregulated in the gametophyte, while the remaining were downregulated including three in the CRM6 clade (Fig. 3C). Interestingly, (67) also identified several CRM3 genes with greater expression in the gametophyte than in the sporophyte, while CRM6-III genes exhibited the opposite pattern. There were clear differences in expression levels in CRM3 and CRM6 genes in *L. microphyllum* gametophyte and sporophyte tissues. Five MADS-box genes also had significantly different expression profiles in *Ceratopteris.* Four of these were downregulated in the gametophyte, including two CRM6 homologs, while an AGL15 homolog was upregulated in mature gametophytes. The differential expression of these MADS-box genes in these two ferns separated by >250 my of evolution suggest a conservation of the pathways integral to gametophyte and sporophyte differentiation and development, though differential expression was not observed in *Adiantum*. In contrast to seed plants where expression of MADS-box genes have specific sporophytic and gametophytic expression patterns (68), MADS-box genes are widely expressed in both life phases in seed-free plants (7, 69), although the expression of genes such as CRM3 and CRM6 may play important roles in tissue differentiation between life stages. Both homeobox (70) and MADS-box genes (71) act as master regulators of plant development in both life phases (72, 73). A comprehensive understanding of the regulation and expression of these genes in gametophyte and sporophyte development will undoubtedly shed light on the generation of morphologically and functionally different life phases from a single genome.

Post-transcriptional modifications, such as RNA splicing, are important processes that contribute to responses to environmental changes, gene regulation, and proteome diversity in plants (74). mRNA from a single gene in complex eukaryotes can be alternatively spliced into different transcripts, or isoforms, and can be classified into four main event types: exon skipping, intron retention, alternative donor, and alternative acceptor (Fig. 3E). Very little is known about splicing in ferns, although some authors have suggested it may play a role in development (e.g., (75), but see (8)). We examined splicing events in the gametophyte and sporophyte to explore the splicing landscape in a homosporous fern. In alignment with data from other plants, the most common splicing event was intron retention (>40% of events, Fig. 3E, Table S11) in both the gametophyte and sporophyte tissue. Previous authors have proposed that intron retention is an important mechanism in plant development and stress responses (e.g., (76)), and could have a role in promoting tissue-specific functions (77). Of the ∼1,700 splicing events identified in each life phase, about a quarter were shared (404 total events); there were 1,363 and 1,302 unique events in the gametophyte and sporophyte, respectively (Fig. 3E, Table S11) and the proportion of each event was approximately equal when compared between tissue types. We compared which genes had unique splicing events in each life stage to the homeobox and MADS-box genes we identified above and manually verified the events. We identified exon skipping in the gametophyte for g22068, a fern-specific HDZIP gene, and intron-retention in the gametophyte for g4788, a MADS-box gene. Whether alternative splicing is important in the context of gametophyte and sporophyte development in ferns requires further study, but our results reveal that there are patterns in both expression and splicing that are unique to each life stage.

### Epigenomic Comparison of Fern Life Phases

DNA methylation is an important mechanism through which eukaryotes regulate gene expression without changing the underlying genetic code (78) and may play a role in fern development (79, 80), but has only begun to be explored in seed-free plants (81). In plants, cytosine methylation is most common at CG sites across the genome, although it is also present in CHG and CHH (where H is A,T, or C) contexts (82). We re-affirmed that the genetic machinery for DNA methylation is largely present in fern genomes (Fig. S9- 11) (6, 7)). Using a phylogenomic approach, we identified homologs of genes involved in *de novo* methylation (*DRM1/2*), the maintenance of methylation (*DRM1/2, CMR1/2, DDM1,* and *MET1*), and demethylation (*ROS1*) that are present across eukaryotes. It is possible that ferns also have homologs of the demethylation gene *DME*, although it was not possible for us to differentiate between *ROS1* and *DME* as their protein products have very similar motifs (78).

To explore the epigenomic landscape of ferns, we generated the first genome-wide methylation data at the single base resolution for a fern gametophyte (pooled immature and mature gametophytes) and compared these data to methylation patterns in the sterile leaf (hereafter ‘sporophyte’ in this section). The genome of both life phases was highly methylated, although the gametophyte had lower methylation (mean 72.63% CpG sites methylated) compared to the sporophyte (93.75%), similar moderate CHG methylation levels (62.70% in the gametophyte, 69.43% in the sporophyte), and low levels of CHH methylation (1.90% in the gametophyte, 1.46% in the sporophyte, Table S12). Both the CG and CHG methylation levels in *Lygodium* are similar to those reported for leaf methylation levels in the tree fern *Alsophila spinulosa* (88.87% CpG, 66.83% CHG), although the CHH methylation level appears to be slightly greater in *Lygodium* (1.68%) compared to *Alsophila* (0.03%) (6). DNA methylation in plants is typically used to silence TEs (83), and given the high density of repeats in fern genomes it is unsurprising to see consistently high DNA methylation levels across the genome (Fig. 4; (6)). Interestingly, very few sites were significantly differentially methylated between the gametophyte and sporophyte; 13,513 CpG sites were hypermethylated in the sporophyte, while just 7,229 CpG sites were hypomethylated in the sporophyte relative to levels in the gametophyte (Fig. 4). The majority of differentially methylated sites were located in intergenic regions (70.34% hypermethylated sites, 64.77% hypomethylated sites), followed by putative promoters (14.08% hypermethylated sites, 29.63% hypomethylated sites). When we examined the putative functions of genes within 10 kb of these differentially methylated sites, the hypomethylated regions were associated with protein kinases (GO:0016773, GO:0004672, GO:0016301). No other functions were enriched in these regions. Sex determination in *Ceratopteris* gametophytes appears to be accompanied by large-scale epigenetic changes (79), although there do not appear to be widespread differences in the epigenome of the gametophytes and sporophytes in *Lygodium*.

**Figure 4.**
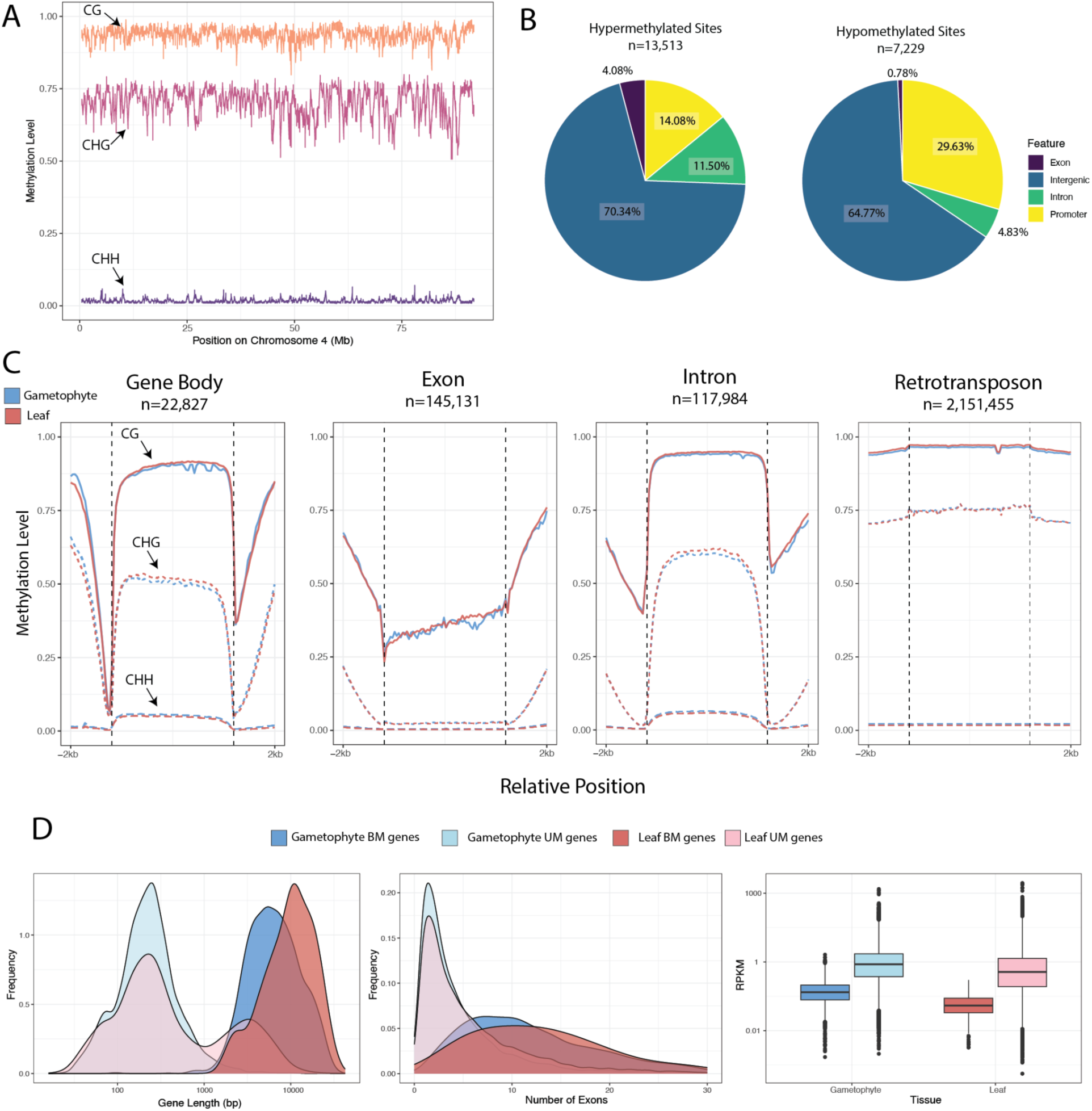
Epigenomic differences in gametophyte and leaf tissues. **A)** Distribution of DNA methylation in CG (orange), CHG (magenta), and CHH (purple) contexts over chromosome 4, as an example, in *Lygodium*. **B)** Classification of regions with differentially methylated sites between gametophyte and leaf tissues. Hypomethylated sites are those that are significantly less methylated in the leaf than the gametophyte. Hypermethylated sites are those that are significantly more methylated in the leaf than the gametophyte. **C)** Distribution of DNA methylation in CG (solid line), CHG (short dashed line), and CHH (long dashed line) contexts over the entire gene body, exons, introns, and retrotransposons for gametophyte (blue). **D)** Plots showing the distribution of gene length, number of genes, and expression level (RPKM) for body-methylated (BM) and unmethylated (UM) genes identified in the gametophyte and leaf tissues.

Gene body methylation (gbM) is particularly interesting to plant biologists as its function is not fully understood and appears to be regulated through a different mechanism than the methylation of TEs (84). Researchers have proposed that gbM may be involved in the stabilization of gene expression, upregulation of gene expression, or preventing TE insertion into vital genes (reviewed by (84)). Recent genomic studies have revealed that gbM is relatively common in ferns (6, 7, 85), but the gametophyte has not been included in these analyses thus far. To explore how gbM is distributed in the life phases of a fern, we compared gbM between gametophyte and sporophyte tissue in *Lygodium.* We identified substantially more body methylated (BM) genes in the gametophyte (2033) compared to the leaf tissue (162), with the opposite trend for unmethylated (UM) genes (10101 in the gametophyte, 16125 in the leaf, Table S13). Nearly all the leaf BM genes also exhibited gbM in the gametophyte (144 shared BM genes), and most UM genes in the gametophyte were also UM in the leaf (9915 shared UM genes). The proportion of gametophyte genes classified as BM (ca. 20%) is similar to that in *Arabidopsis* (ca. 27% of genes (86)) and across angiosperms (ca. 18% (87)), whereas the proportion of BM genes in the leaf is substantially lower (ca. 1%). Such a vast difference could be due to higher background methylation levels in the leaf tissues, making it difficult to pull out significant gbM using the binomial approach used here. Alternatively, differences in the number of BM genes could be due to extensive changes in DNA methylation in the life cycle, which are known to occur in *Marchantia polymorpha* (88) and *Arabidopsis thaliana* (89).

To determine whether BM genes are enriched for functional categories, we performed a GO enrichment test of the BM genes independently in each tissue relative to the total genome annotation. Housekeeping genes, which have constitutive and moderate expression, generally exhibit gbM in invertebrates (90, 91). Our results largely agree with these findings (Table S14): both BM genes in the gametophyte and leaf were enriched for processes related to lipid metabolism (e.g., GO0042157, GO:0042158), while the gametophyte BM genes were also enriched for protein localization (GO0033365, GO:0008104) and hydrolase activity (e.g., GO:0016787, GO:0016811). Interestingly, in both gametophyte and leaf tissues BM genes had lower expression levels (measured by RPKM within samples) compared to UM genes (Fig. 4D), which contrasts with findings in *Arabidopsis* where BM genes were moderately expressed and UM genes had more variable expression profiles (86). The general characteristics of the BM genes, however, were similar to what is known in angiosperms. Consistent with the expectation that gbM is associated with stabilizing expression (86), gene length in BM genes were significantly greater than UM genes in both the gametophyte (exact permutation test estimated by Monte Carlo with 999 permutations *P =* 0.002) and leaf tissues (*P =* 0.002); the leaf BM genes were significantly longer than the gametophyte BM genes on average (*P* = 0.002; Fig 4D). We also observed that BM genes had a greater number of exons in the gametophyte (*P=*0.002) and leaf (*P =* 0.002), but there was not a difference between the gametophyte and leaf BM genes (*P* = 0.396; Fig. 4D).

Our analysis of the first methylome of a fern gametophyte has revealed similar patterns of methylation in the haploid gametophyte life phase compared to the sporophyte. A slightly lower global methylation level in the gametophyte did not correlate with the very few differentially methylated sites between life phases. Our approach using pooled gametophyte tissues may swamp out the signal of tissue-specific methylation patterns in the gametangia and gametes that was observed in *Marchantia* (*88*).

Epigenetic marks are an important mechanism through which plants can regulate gene expression (92) and may play a vital role in the differentiation of gametophyte and sporophyte genetic programs, and further investigation will be required to untangle these processes in plants.

### Phase-Specific Cold Tolerance Highlights Resilience of the Gametophyte

Freezing is widely predicted to be the most likely abiotic factor limiting range expansion of *L. microphyllum* (93). We used an ecological niche model to identify specific bioclimatic variables with high permutation importance in describing the distribution of this species, which were isothermality (bio3), temperature seasonality (bio4), and mean temperature of the warmest quarter (bio10, Table S15). Given the importance of temperature in the niche suitability of this species, *Lygodium microphyllum* is likely to experience a northward range shift as the climate continues to change (Fig. 5; (35)), and the tolerance of the gametophyte and sporophyte life stages to these limiting variables will determine the spread of its invasion.

**Figure 5.**
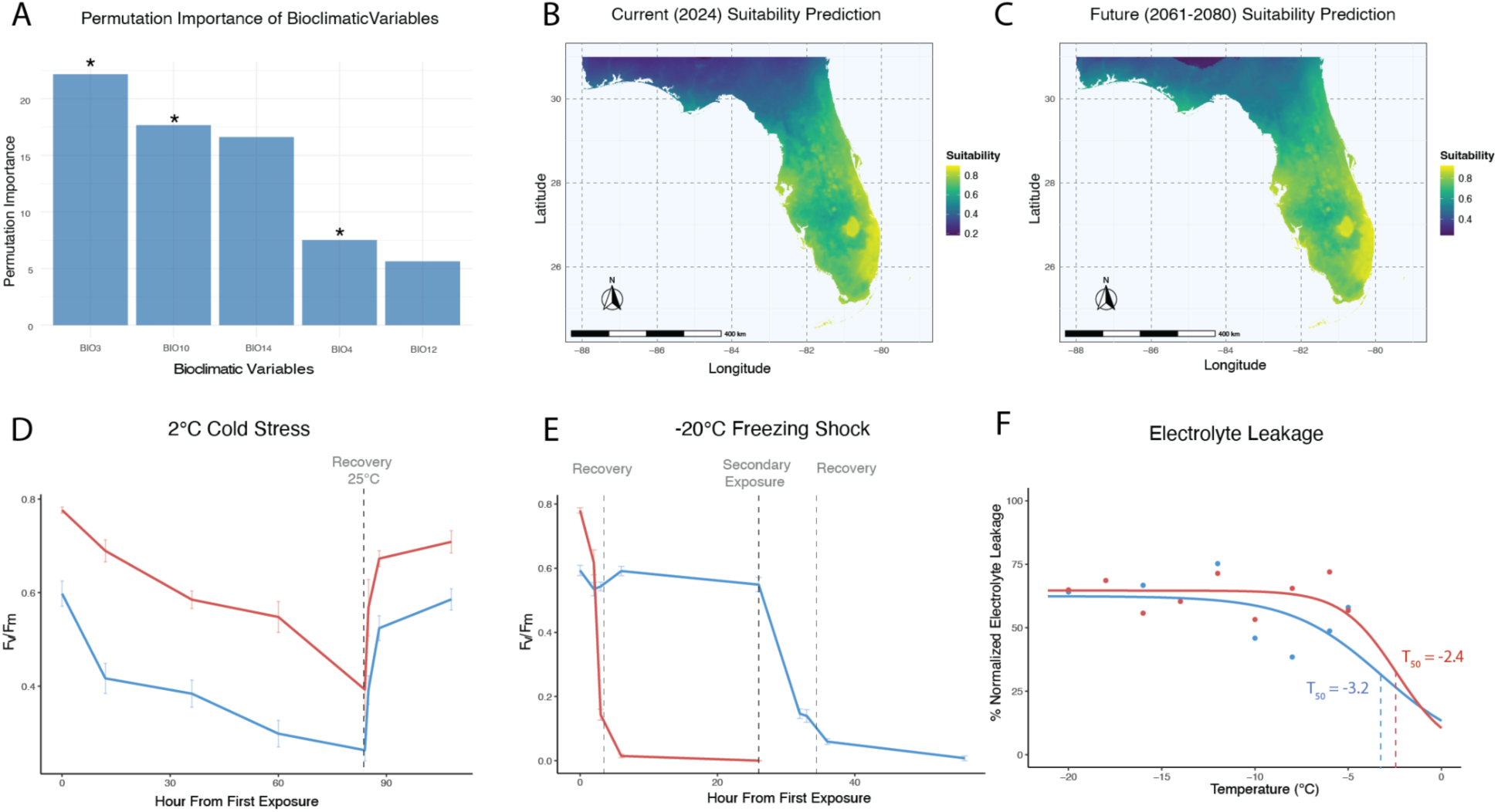
Differential physiological responses to cold stress and freezing shock. **A)** Permutation importance of climatic variables >5% within the niche model for *Lygodium microphyllum*. Variables associated with temperature (bio3, bio10, and bio4) are denoted with an asterisk. Predicted suitability of *L. microphyllum* during the present (**B**) and projected onto a future climate scenario **(C)** for 2061-2080. A clear northward expansion in suitability is evident in the future climate scenario. **D)** Gametophyte and leaf stress responses to exposure to 2°C temperatures for 84 hours and recovery at 25°C for 24 hours as measured by changes in the maximum quantum yield of photosystem II (Fv/Fm). **E)** Responses of gametophyte and leaf to freezing shock (exposure to -20°C) for 2 hours and recovery at 25°C for 24 hours as measured by Fv/Fm. Gametophytes were exposed to a secondary, longer period (4 hours) after recovery. **F)** Electrolyte leakage of gametophytes and leaf tissue, highlighting a slightly greater ability of the gametophyte to withstand freezing damage.

To understand how expression of the genome might shape physiological responses to limiting environmental conditions at each life stage, we measured physiological responses to cooling and freezing stress in both the gametophyte and sporophyte, and compared gene expression in these tissues. Several large-scale cellular changes are expected to occur during freezing stress. Most importantly, ice will form in the apoplast, disrupting the cell membrane, causing cell dehydration (94) and inducing osmotic shock (95). These changes directly impact the efficiency of the photosynthetic machinery, particularly photosystem II (96). Therefore, we first compared the maximum quantum yield of photosystem II (F_v_/F_m_) of sporophyte and gametophyte samples during the response to and recovery from prolonged cold stress at 2°C (Tables S16-17). As a measure of chlorophyll fluorescence, F_v_/F_m_ can be used as a proxy for plant stress, where values closer to zero represent more stressed plants. There was a significant impact of life stage and duration of cooling at 2°C on the efficiency of photosystem II (linear regression, F_3,118_ = 78.32, *P* = 6.42 x 10^-14^, adjusted R^2^=0.66). Sporophytes had, on average, greater F_v_/F_m_ values than gametophytes during exposure to cold temperatures (0.23 higher F_v_/F_m_ values, *P* = 2.48 x 10^-10^) and for every 10 hours of cooling stress F_v_/F_m_ decreased by 0.03 independent of life stage. While there was a significant effect of duration of recovery on F_v_/F_m_ (i.e., when plants were removed from 2°C and placed at 25°C; log-linear regression, F_4,99_=34.74, *P* = 2.37 x 10^-9^, adjusted R^2^=0.58), there was not a significant difference in the recovery of sporophytes and gametophytes (*P* = 0.26; Fig. 5).

When exposed to short-term freezing stress (2 hours at -20°C), there was a significant impact of freezing and life stage on measures of F_v_/F_m_ (F_3,66_=15.74, *P* = 7.95 x 10^-8^, adjusted R^2^ = 0.39), including a significant faster decrease in F_v_/F_m_ by 0.05 per hour in sporophytes compared to gametophytes at -20°C (*P* = 0.04). Fronds were unable to recover from 2 hours of freezing stress (F_3,101_=250.81, *P* < 2.2 x 10^-16^, adjusted R^2^ = 0.88). Gametophytes returned to pre-freezing F_v_/F_m_ values within 24 hours, but secondary exposure to -20°C temperature for 6 hours resulted in death of the gametophytes. While (34) observed a 7-fold decrease in survival of *L. microphyllum* gametophytes after just 15 minutes at -2.2°C, we found that gametophytes were able to survive far longer durations of freezing stress. Gametophyte cell membranes were also able to withstand slightly lower freezing temperatures (gametophyte T_50_ = -3.2°C, sporophyte T_50_ = -2.4°C, Table S18), although a model including life stage did not improve model fit (ΔBIC =11.5). Importantly, it is well-documented that while fronds of *Lygodium* may die back during freezing events, the rhizome is typically tolerant and will resprout (97). We observed a similar phenomenon, where new fronds emerged within weeks of the freezing shock stress from the rhizome.

### Molecular Pathways Used in Response to Freezing Shock are Tissue-Specific

Given the vastly different physiological responses of the sporophyte and gametophyte to freezing stress, we assessed transcriptomic changes associated with the leaf, gametophyte, and root tissues when exposed to freezing shock. The overall responses varied between tissue types: the leaf had a large number of differentially expressed genes in response to freezing (DEGs, 3903, 19.5% of the total), both the gametophyte (646, 3.3%) and root (415, 2.1%) had very few DEGs (Fig. S12), suggesting that there is a massive transcriptomic change in the leaf but fewer changes in the gametophyte and root in response to freezing. We discuss the changes most striking pathways involved in responses to freezing below (see Table S19, Figs. S13-17).

The remobilization of starch has been implicated in successful abiotic stress responses in several plant species (reviewed by (98)). Interestingly, the gametophyte had increased expression levels of genes in pathways enriched for metabolism, particularly in starch and sucrose metabolism (3.16 F.E., adjusted *P* = 0.0127; Fig. 6a), while the same pathway was enriched in the downregulated genes of the leaf (1.96 F.E., adjusted *P* = 0.0454; Fig. 6a). β-amylases, enzymes that act to release maltose that may act as an osmoprotectant under increased osmotic stress (99, 100), were enriched in these pathways and upregulated in the gametophyte but downregulated in the leaf. The gametophyte also showed enhanced expression leading to the accumulation of trehalose, a small sugar known to improve cold tolerance through several molecular mechanisms including increasing photosynthetic efficiency and ROS scavenging (101). Photosynthetic pathways (e.g., ath00195, 4.49 F.E., adjusted *P* = 0.0489) and proteins (e.g., ath00196, 18.84 F.E., adjusted *P* = 3.12 x 10^-5^) were also enriched in upregulated genes in the gametophyte, supporting the heightened efficiency of photosystem II observed during freezing stress in our physiological experiment (Figs. 5, 6).

**Figure 6.**
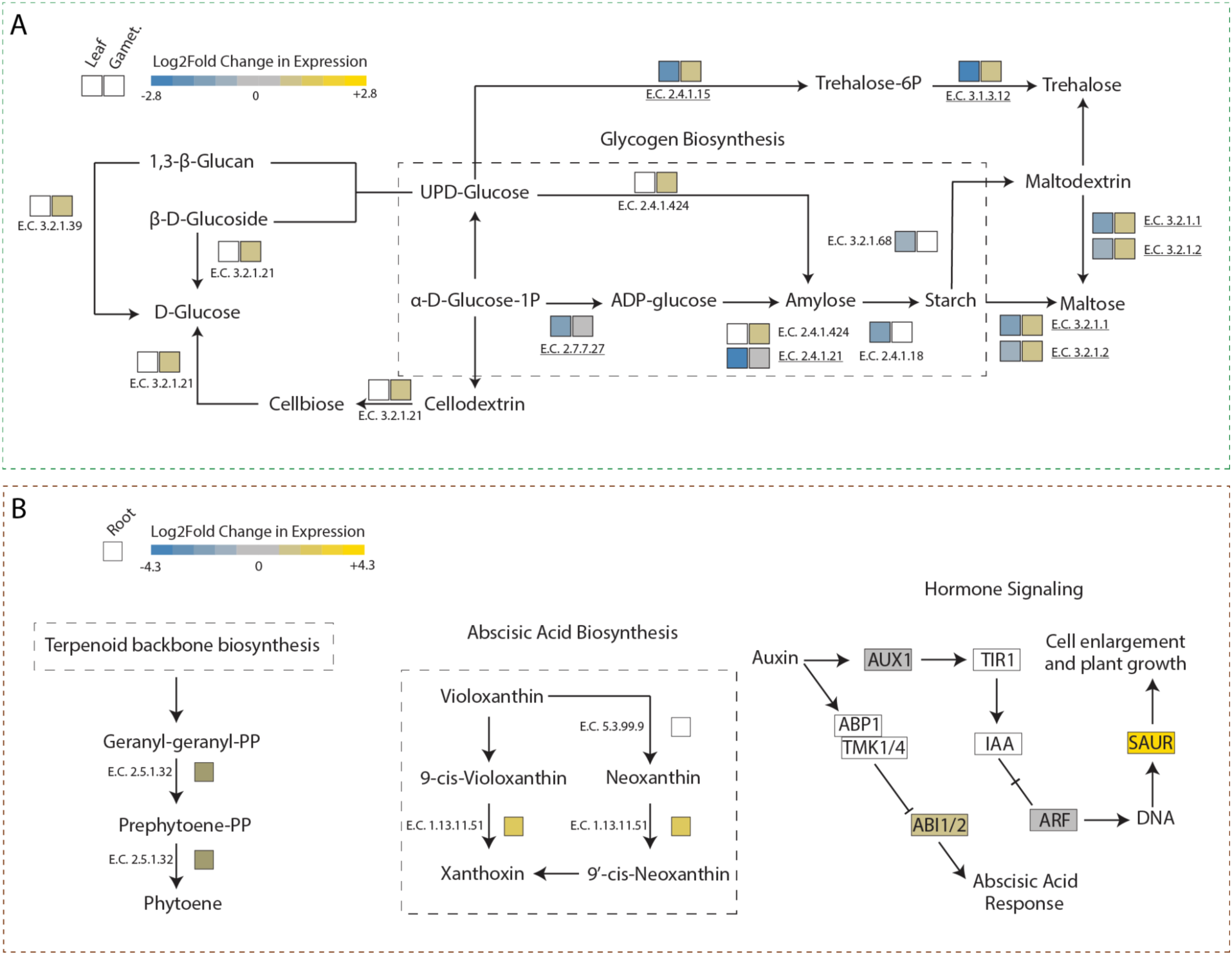
Tissue-specific molecular pathways in response to freezing shock. **A)** Subset of the starch and sucrose metabolism pathway, which was enriched in downregulated genes in the leaf and upregulated genes in the gametophyte as a response to freezing shock. Log2foldchange expression levels of genes involved in enzymatic reactions are listed at each reaction. **B)** Subsets of the carotenoid biosynthesis and hormone signaling pathways that were enriched in upregulated genes in the root as a response to freezing shock. Log2foldchange expression levels of genes involved in enzymatic reactions or hormone-associated genes are listed at each reaction.

In contrast, the root tissue gene sets were not enriched for metabolic pathways (Fig. 6b). Rather, upregulated genes were enriched for the carotenoid biosynthesis pathway (4.23 F.E., adjusted *P* = 2.7 x 10^-5^) and plant hormone signal transduction (11.91 F.E., adjusted *P* = 0.027; Fig. 6b). Carotenoids, and molecules derived from them such as abscisic acid (ABA), are known to play important roles in abiotic stress responses in plants as free-radical scavengers and in regulating and directing plant architecture (102). Studies in angiosperms have highlighted heightened expression of carotenoid biosynthetic enzymes in roots under salt stress, but not shoots (103, 104). Our results agree that the upregulation of carotenoid biosynthesis-related genes is tissue-specific and concentrated in root tissue.

## Conclusion

In this study, we generated a chromosome-scale reference genome of the highly successful invasive vining fern *Lygodium microphyllum* and explored the transcriptomic, epigenomic, and physiological differences between haploid gametophyte and diploid sporophyte life phases. We found that there are fine-scale transcriptomic differences in the regulation of developmental genes and splicing landscape that may explain, at least partially, how a single genome can produce two vastly different life stages. Using the first genome-wide methylation data for a fern gametophyte, we showed that there was remarkable similarity in the epigenomes of gametophytes and sporophytes. By pairing physiological metrics of stress and RNASeq, we were able to demonstrate that the different life stages of this fern likely leverage their shared genome via different strategies to respond to environmental changes. In the gametophyte, which showed heightened freezing tolerance, pathways involved in the accumulation of soluble sugars were activated, while the same pathways were downregulated in the leaf, which quickly succumbed to freezing shock. Our results highlight the tolerance of *L. microphyllum* gametophytes to freezing and suggest that the gametophyte may play a pivotal role in the establishment dynamics and range expansion of *L. microphyllum*, especially as the climate continues to warm. Given that many organisms exhibit functional variation between life stages, determining their underlying biology at multiple scales, from genome to phenotype, across life phases, is vital to developing a thorough understanding of organismal biology.

## MATERIALS AND METHODS

Living plant material (sterile leaf tissue) was collected in Taiwan from an individual currently maintained at the Dr. Cecilia Koo Botanic Conservation Center (voucher Lu 30898 available at the herbarium TAIF, collected from Mt. Tanfeng, Beitou District, Taipei City, Taiwan). For genome assembly, we used a combination of long-read Oxford Nanopore (∼37x), Illumina short-read (∼60x), and chromosome conformation capture (Dovetail’s Omni-C; ∼73x) sequencing technologies. A contig-level assembly was generated with error-corrected Nanopore reads >5kb using HERRO v0.1.0 (105) and hifiasm v0.19.9 (106). The assembly was scaffolded with the Omni-C data using juicer v2.0 (107), YaHS v1.2 (108) and Juice Box Assembly Tools v2.17.00 (109). We used both reference protein and newly generated RNAseq data from nine tissue types to annotate the genome with the BRAKER3 v3.0.6 pipeline (110). Functional annotations of the resulting gene models were assigned with InterProScan v5.68-100.0 (111) and eggNOG Mapper v2.1.12 (112). Analysis of differential gene expression was performed with DESeq2 v1.44.0 (113) and alternative splicing events identified using a modified pipeline from (114, 115). MethylSeq data were mapped to the genome with bwa-meth v0.2.7 (https://github.com/brentp/bwa-meth) and analyzed with MethylDackel v0.6.1 (https://github.com/dpryan79/MethylDackel) and methylKit v1.18.0 (116). To compare gene body methylation (gbM) between gametophyte and sporophyte tissues, we used the probabilistic approach described by (86). Details on the materials and methods used in this study can be found in the Supplemental Materials and Methods.

## DATA, MATERIALS, AND SOFTWARE AVAILABILITY

Sequence data have been deposited in NCBI SRA under BioProject XXXXXXX and from 10KP at https://db.cngb.org/search/organism/148566/. The genome assembly is available at NCBI under accession XXXXXXXXXXXX. Genome assembly and annotations are also available at https://github.com/jessiepelosi/LygodiumGenome and FernBase (fernbase.org). All code used in this study is available at https://github.com/jessiepelosi/LygodiumGenome.

## ACKNOWLEDGEMENTS

We are grateful to Nathan Backenstose, Katherine Eaton, C. George Glen, Christopher Kreig, Qinyin Ling, Poppy Northing, Kasey Pham, Weston Testo, Eddie Watkins, and Bethany Zumwalde for comments, suggestions, and advice on this project; Katherine Eaton and the Bernal lab at Auburn University and Chris Dervinis at the University of Florida for their assistance with HMW DNA extractions; Charlie Bear, Bill Hammond, Grace John, Nick Keiser, and Zepeng Yao for use of equipment and lab space. We thank the 10,000 Plant Genomes Initiative, Sunil Kumar Sahu, and Gane Ka-Shu Wong for providing short read data they generated as part of 10KP. The authors acknowledge University of Florida Research Computing for providing computational resources and services that have supported this research.

Funding for this project was provided in part by the Florida Invasive Species Council Julia Morton Grant, University of Florida Grinter Fellowship, University of Florida Department of Biology Mildred Mason Griffith Botany Grant and Michael May Dissertation Fellowship, Botanical Society of America J.S. Karling Graduate Student Research Award, and USDA NIFA Postdoctoral Fellowship #2024-67012-43394 to J.A.P. and NSF IOS #1754911/2310485 to E.B.S. A.J.D. was supported by USDA NIFA Predoctoral Fellowship #2024-67011-43011 and K.M.D. was supported by USDA NIFA award #2023-67013-40169. Movement of plant parts was conducted in accordance with USDA permit P526P-21-02509.

