## Supplemental Information for "The genome of the vining fern *Lygodium microphyllum* highlights genomic and functional differences between life phases of an invasive plant"

#### SUPPLEMENTAL MATERIAL

**Includes:**

Supplemental Materials and Methods
Supplemental Figures 1-17

### SUPPLEMENTAL MATERIALS AND METHODS

#### Plant Material and Sampling

Living plant material of an individual *Lygodium microphyllum* plant (K056944) is maintained at the Dr. Cecilia Koo Botanic Conservation Center (KBCC). A voucher specimen is deposited at the Herbarium of Taiwan Forestry Research Institute (TAIF; specimen Lu 30898, collected from Mt. Tanfeng, Beitou District, Taipei City, Taiwan). Spores from the KBCC individual were sown on Bold's media (1) with Nitsch's micronutrients (2) and gametophytes were grown at an average temperature of 23.5°C on a 12:12 light:dark schedule (average light intensity 30.4 $\mu mol/m^2/s$). All sporophytes were produced by sporophytic selfing (a gametophytically selfed sporophyte was produced but had substantial developmental stunting and did not grow past a few cm tall). The resulting sporophytes were the primary source for genome and transcriptome sequencing.

#### Genome Sequencing

Short read whole genome sequencing data were generated by the 10,000 Plant Genomes Project (3). A total of 751.8M 2x100 reads (75.1 Gb; 15x coverage) were sequenced on a BGISEQ-500 following the standard protocols of the 10KP project. DNA for short-read sequencing was extracted using the Qiagen Plant DNEasy kit following the manufacturer's protocol. Library preparation and Illumina sequencing (2x150bp) were completed at NovoGene on an Illumina HiSeq 2500; 1.5B genomic reads (230.7 Gb) were generated. In total, we generated ~60x coverage (305.8Gb) of short read data.

HMW DNA was extracted from sterile leaf using a nuclei isolation (4) with woody plant buffer (5), SDS-based extraction (6), and high salt clean-up (7). Prior to extraction, plants were dark-treated for at least 48 hours to reduce photosynthate and other organic compounds during the extraction process. Long-read sequencing was performed using two flow cells on the PromethIon-24 at the University of Florida Interdisciplinary Center for Biotechnology Research (ICBR) NextGen Sequencing Core (RRID:SCR_019152) and three version R10 flow cells in-lab at the University at Buffalo. For PromethIon runs at ICBR, we sheared DNA to an average size of 20kb using a Covaris g-tube and removed fragments <10kb with the PacBio SRE XS eliminator kit. One flow cell was run following size selection with a PacBio SRE XL eliminator kit at the University at Buffalo. A total of 19.7M reads (191.68Gb) with an N50 of 14.4kb (individual library read N50s ranged from 11kb - 26kb) were generated from the Nanopore runs to an approximate depth of coverage of 37.6x (based on an estimated genome size of 5.1Gb). All ONT data were basecalled using the super accuracy model in real-time in MinKnow v23.04.5 with Guppy v6.5.7 or were subsequently basecalled with Dorado v0.4.2 (https://github.com/nanoporetech/dorado) prior to assembly.

To generate chromatin conformation data, we first isolated nuclei following the protocol from (8). Approximately 1g of sterile leaf tissue was ground in liquid nitrogen for nuclei isolation; we ensured that intact nuclei were obtained by staining with DAPI and examining under a fluorescent scope. The pelleted nuclei were washed with PBS, snap frozen in liquid nitrogen, and then used as the input into the Omni-C Mammalian Cell protocol following the manufacturer's protocol (Dovetail Genomics, CA, USA) using a 1:10 dilution of the nuclease enzyme mix and 25-30 min digestion. Three libraries were generated from the digestions following the manufacturer's protocol and sequenced on one lane of Illumina NovaSeqX 10B at the UF ICBR. Each library was sequenced to approximately 350M read pairs for a total of 351.3Gb of data, approximately 73.2x coverage.

**Genome Assembly**

Genome size and heterozygosity were first assessed by generating a 61-mer frequency distribution with Jellyfish v2.3.0 (9) and analyzed in GenomeScope v2.0 (10) and a custom R script. We used HERRO v0.1.0 (11) to correct ONT reads >5kb in length. The corrected reads were assembled with hifiasm v0.19.9 (12). We verified the quality of the assembly by mapping genomic Illumina short reads with bwa v0.7.17 (13). OmniC data were mapped to the contig-level assembly and filtered using the juicer v2.0 (14) pipeline. Contigs were then scaffolded with YaHS v1.2 (15) and manually adjusted as needed with Juice Box Assembly Tools v2.17.00 (16). The completeness of the genome assembly was assessed with compleasm v0.2.2 (17) with the Viridiplantae odb10 database from BUSCO (18). We used tidk v0.2.14 (19) to identify telomeric repeats and visualize their location in the scaffolded genome assembly. NCBI's foreign contaminant filter (20) was used to remove any possible non-fern sequences (adaptor and prokaryotic contamination). A total of 69 unplaced contigs totalling 2.8 Mb were removed from the final assembly. All data analysis was conducted on the University of Florida's HiPerGator computing cluster.

**Transcriptome Sequencing**

Tissues for transcriptome sequencing were flash frozen in liquid nitrogen and stored at -80°C until extraction. RNA was extracted from 2 to 3 biological replicates of sterile leaf, fertile leaf, elongating leaf, rhizome, rachis, and fiddleheads using a modified CTAB extraction (21), purification with the Spectrum Plant Total RNA Kit or Zymo RNA Clean and Concentrator and on-column DNAse treatment. mRNA Illumina libraries were constructed using this RNA and sequenced at NovoGene to an average depth of 20 million 2x150 read pairs per library. Additional libraries were generated from the freezing experiment (see below; sterile leaf, pooled gametophytes, root/rhizome) and sequenced by Novogene as above with stranded RNASeq libraries. Two additional sterile leaf libraries were generated and sequenced by 10KP (3) on a BGISEQ-500 following standard protocols of the project.

**Plastome Assembly and Annotation**

The plastid genome was assembled with a random 1Gb subset of BGI reads generated by 10KP with Novoplasty v4.2 (22) using the default parameters with the *Arabidopsis* rbcL sequence as the seed. The plastome was annotated with PGA v1.0 (23).

**Genome Annotation**

A species-specific repeat library was constructed with RepeatModeler (24) and repetitive elements were masked with RepeatMasker v4.0.5 (25). Transcript-based evidence was generated by mapping RNASeq reads to the masked genome assembly with hisat2 v20230203 (26)). Protein-based evidence was derived from 16 proteomes: *Chlamydomonas reinhardtii* (27)*, Anthoceros agrestis* Bonn (28), *Marchantia polymorpha (29), Physcomitrum patens (30), Selaginella moellendorffii (31), Azolla filiculoides (32), Salvinia cucullata (32), Marsilea vestita (33), Ceratopteris richardii (34), Adiantum capillus-veneris (35), Alsophila spinulosa (36), Amborella trichopoda (37), Oryza sativa (38), Zea mays (39)*, *Populus trichocarpa (40),* and *Arabidopsis thaliana (41)*. RNASeq and protein evidence were used to generate gene models within BRAKER 3(42), which relies on AUGUSTUS (43,44), DIAMOND (45), GeneMark-ETP (46), GFF utilities (47), SPLAN (48,49), and StringTie (50). The completeness of the resulting gene prediction was determined with BUSCO v5.3.0 (18) with the Viridiplantae odb10 dataset in proteome mode and Omark web browser (51). Statistics for the gene annotation were generated with AGAT v0.4.0 (52). We used InterProScan v5.68-100.0 (53) and eggNOG Mapper v2.1.12 (54) with the Viridiplantae eggNOG database v5.0.2 (55) to assign putative functions to our gene models.

**DNA Methylation**

DNA was extracted from three pooled (ca. 3-5) gametophyte and three young sterile leaf samples of *Lygodium microphyllum* following a modified CTAB extraction and purification with the Qiagen Plant DNeasy kit. Methyl-seq libraries were constructed using the NEBNext Enzymatic Methyl-seq Kit and sequenced to an average depth of 10x (8.8-13.7x) per library on the Illumina NovaSeq Sp4 at the UF ICBR. Read quality was assessed with FASTQC, adapters trimmed and the first five bp removed with Trimmomatic v0.39 (56), and aligned with bwa-meth v0.2.7 (https://github.com/brentp/bwa-meth). PCR duplicates were removed with Picard in GATK v4.3.0.0 (57) and methylation calls performed in MethylDackel v0.6.1 (https://github.com/dpryan79/MethylDackel). Differential methylation at the base pair resolution between gametophyte and sporophyte tissues was called with methylKit v1.18.0 (58) in R v4.1.2 (59) with using Fisher's exact test, requiring at least 5x coverage. Methylation was visualized with ViewBS v0.1.10 (60) and custom R scripts.

To compare gene body methylation (gbM) between gametophyte and sporophyte tissues, we used the probabilistic approach described by (61) to determine which genes were significantly methylated in the CpG context. We removed bases from the analysis that had less than 2x coverage and genes that had fewer than 20 total cytosines. (61) used a binomial distribution to calculate a one-sided p-value (e.g., $P_{CG}$) for each gene compared to the genome background. In R:

$$\text{binom.test}(x=mcg,\ n=ncg,\ p=G_{CX},\ \text{alternative} = \text{“greater”})$$

where *mcg* is the number of methylated cytosines, *ncg* is the total number of cytosines, $G_{CX}$ is the average proportion of methylation in a given context. Since we had six different libraries with varying global genomic mCG levels, we set the genome background independently for each library. We then corrected $P_{CG}$ values for multiple comparisons using the Bonferroni method and classified genes as body-methylated (BM) if the adjusted $P$-value < 0.05, intermediately methylated (IM) if 0.05 < adjusted $P$-value < 0.95, and unmethylated if $P$-value > 0.95. We performed the binomial test for each context (CG, CHG, and CHH) independently. Given that true BM genes only have significantly greater methylation in the CG context (62) and to be consistent with previous studies (61,63,64), genes with $P_{CHG}$ and $P_{CHH}$ < 0.05 were removed from the list of BM genes. We only retained BM and UM genes that were shared between all three libraries for each tissue type to give a conservative set of genes. We compared the distributions of gene lengths (including exons and introns, from start to stop codon) and the number of genes between BM and UM genes in each tissue type using exact permutation tests estimated by Monte Carlo with 999 permutations in the R package perm 1.0-0.4 (65). We used GO term enrichment, performed with TopGO v2.56.0 (66), to determine if there were functional categories enriched in BM and UM genes relative to the total genome annotation. In order to compare expression levels of BM and UM genes, we used edgeR v4.2.2 (67) to calculate RPKM values for BM and UM within each library. RPKM is a normalization method that accounts for sequencing depth and gene length, but cannot be used to compare expression between samples.

**Phylogenomics**

Proteome and coding sequences were downloaded for 19 species of plants including some of the species used in genome annotation, in addition to *Diphasiastrum complanatum* v3.1 *(68)* and *Isoetes taiwanensis (69)* and excluding *Selaginlla moellendorfii.* We inferred the phylogeny of land plants by extracting 213 genes identified as low-copy by GoFlag (70). We identified these genes using tblastx (71) against the reference loci from GoFlag and retained only the longest isoform from each species. Sequences were aligned with the codon-aware alignment program MACSE v2.0.4 (72) and gappy sites were removed with trimAl v1.2 (73) by retaining sites that contained at least 50% of tips. The best-fitting substitution model was determined with ModelFinderPlus (74) and maximum likelihood (ML) gene trees were constructed from the nucleotide alignments using IQTREE2 v2.1.2 (75) with 1000 ultrafast bootstraps (76). A multi-species coalescent tree was constructed from the resulting gene trees with ASTRAL v5.15.5 (77). We generated an ML species tree based using the ASTRAL topology as a constraint and the concatenated matrix of 213 genes. This tree was used to estimate divergence times among major lineages of land plants with treePL v20150305 (78). We used 1000 ultrafast bootstraps to account for uncertainty in the dataset and ran treePL on these bootstrap trees and summarized the supports using scripts from https://github.com/sunray1/treepl. Eight calibrations points were used based on the estimated minimum and maximum ages from (79): crown Viridiplantae (469-1891 Mya), crown Embryophyta (469-515.5 Mya), crown Marchantiophyta + Bryophyta (405.0-515.5 Mya), crown Tracheophyta (420.7-451 Mya), crown Lycopodiophyta (392.1-451 Mya), crown Euphyllophyta (385.571-451 Mya), crown Spermatophyta (308.14-365.629 Mya), and crown Angiospermae (125-247.2 Mya).

**Alternative Splicing and Differential Gene Expression between Life Phases**

RNA sequence data from control young, sterile leaf and gametophyte tissues (three replicates each) were used to compare the transcriptomic landscape of the gametophyte and sporophyte life stages since these tissues are functionally the most similar. Hisat2 v20230203 (26) was used to map RNASeq reads to the genome and read counts generated with HTSeq v2.0.3 (80) with parameters: -m union -t mRNA -s reverse). Differentially expressed genes were identified in DESeq2 v1.44.0 (81) filtering for LogFoldChange > 2 and adjusted p-value < 0.05. Analysis of enrichment of KEGG (82) pathways in upregulated and downregulated gene sets was performed with clusterProfiler v4.12.6 (83) and GO term enrichment was performed with TopGO v2.56.0 (66).

To compare our results to other ferns, we downloaded RNASeq data from (34,35) that most closely matched the tissue types that we generated. From Fang et al. (2022) we used the "transEG" (SRR11613609, SRR11613614, SRR11613662) and "transUL" (SRR11613625, SRR11613636, SRR11613647) datasets which are the embryonic gametophyte and unfurled leaf tissues, respectively. From (34) we used sterile leaf (SRR17380274, SRR17380275, SRR17380276) and mature gametophyte libraries (SRR17275295, SRR17275322, SRR17275323, SRR17275324). Data analysis of these libraries followed the pipeline used for *Lygodium* above.

We used OrthoFinder v2.5.5 (84,85) to identify MADS-Box and homeobox gene families, as classified by TAIR (86) and (87). Orthogroups containing *Arabidopsis* homologs of interest were extracted and proteins aligned with MAFFT v7.5.20 with the integration of structural information with Database of Aligned Structural Homologs (DASH) (88). Sites with less than 20% occupancy were removed, leaving conserved protein domains. Individual orthogroups were aligned with MAFFT v7.5.20 and sites with less than 50% occupancy were removed and gene trees were constructed as below. The best-fitting substitution model was determined with ModelFinderPlus (74) and the maximum likelihood gene tree was constructed from the protein alignment using IQTREE2 v2.1.2 (75) with 1000 ultrafast bootstraps (76).

To identify alternative splicing events between the gametophyte and sporophyte, we used a modified version of the pipeline described in (89,90). RNASeq libraries for control gametophyte and leaf tissues were merged with samtools v1.20 (91,92). Transcripts were generated for the three the gametophyte and sterile leaf merged BAMs using Stringtie v2.2.3 (93) with default settings (minimum isoform fraction = 0.01 and minimum FPKM value = 1.0). We then used PASA v2.4.1 and v2.5.3 (94) to identify all alternative splicing events with minimap2 v2.28 (95) and filtered events to only include those with more than two reads supporting each junction and reclassified events according to (89). Shared and unique events were identified and compared to gene families of interest (homeobox and MADS-box genes) in R.

**Ecological Niche Modeling**

A species distribution model was constructed using Maxent v3.4.3 (96) through the R package dismo v1.3-14 (97). The present-day model was generated using all 19 WorldClim variables at 30s resolution (98). The content of Antaractica was removed from the model. Collinearity between environmental variables did not strongly affect model output and therefore all 19 variables were retained. Cleaned occurrence records for *Lygodium microphyllum* were retrieved from (99), and a bias layer was generated for Lygodiaceae. The model achieved an AUC of 0.944 based on the training data and bioclimatic variable contribution and permutation importance can be found in Table S15. The model of the current distribution was projected onto a future climate scenario to predict future suitability. For the future climate scenario, we used the MIROC6 model (100) with shared socio-economic pathway SSP 370 at 30s spatial resolution for the 2061-2080 time period.

**Cold/Freezing Stress Experiments**

*Experiment 1: Cooling Stress*

Sporophytes and gametophytes were placed in a Percival I-36LL growth chamber (Percival Scientific, IA, USA) on a 12:12 light:dark schedule at 25°C and allowed to adjust to these conditions for two weeks. Prior to ramping down temperature, control measurements of $F_v/F_m$ were taken with a MiniPam II (Heinz Walz, Effeltrich, Germany). Temperature was ramped down over 8 hours from 25°C to 2°C and then maintained at 2°C in the growth chamber on the same 12:12 light:dark schedule. Measurements of $F_v/F_m$ were taken at 12-, 36-, 60-, and 84-hour timepoints during cold stress at 2°C and 1, 4, and 24 hours of recovery (25°C) using the MiniPam II. Samples were incubated in the dark at the treatment temperature for at least 15 minutes prior to measurements. The fluorometer was maintained at 60° and 45° angles for sporophyte and gametophyte samples, respectively, and at a constant distance of 3 mm from the sample.

*Experiment 2: Freezing Stress*

Sporophytes and gametophytes were incubated as above for the control measurements. Plants were then moved to a -20°C freezer for 2 hours and then allowed to recover at 25°C for 24 hours. Fluorescence measurements were taken immediately after removal from the freezer, and 1, 4, and 24 hours during recovery. Gametophytes were subjected to a repeated freezing event after 24 hours for an additional 6 hours at -20°C; measurements were taken as above. The experiment was repeated with a second set of sporophyte and gametophyte samples without taking $F_v/F_m$ measurements. Tissues were immediately frozen in liquid nitrogen at the control 25°C and after 2 hours at -20°C and RNA was extracted from leaf, root/rhizome, and gametophyte tissues. Hisat2 v20230203 (26) was used to map RNASeq reads to the genome and read counts generated with HTSeq v2.0.3 (80) with parameters: -m union -t mRNA -s reverse). Differentially expressed genes were identified in DESeq2 v1.44.0 (81) filtering for LogFoldChange > 0 and adjusted p-value < 0.05. Genes related to CBF regulation and the AP/ERF and WRKY families were identified by blasting the *Lygodium* proteome against *Arabidopsis thaliana* gene models with BLASTP (71). Analysis of enrichment of KEGG (82) pathways in upregulated and downregulated gene sets in each tissue type was performed with clusterProfiler v4.12.6 (83)

*Experiment 3: Electrolyte Leakage*

Electrolyte leakage was assessed for both sporophyte and gametophyte tissue at multiple freezing temperatures following a protocol modified from (101,102). Briefly, freezing was induced in leaf discs from sporophytes or whole gametophytes in 5 mL of ultrapure water. Samples were ratcheted at twodegree intervals from 1°C to -20°C and were kept at the treatment temperature for 30 minutes before being removed or continuing to the next temperature. Samples were then placed on ice and thawed overnight and then shaken at 150 rpm for 16 hours at room temperature. Conductivity measurements were taken with an Orion Star A325 conductivity/pH meter (Thermo Scientific, Waltham, MA, USA). Samples were autoclaved to cause complete cell lysis, shaken for 16 hours at room temperature, and conductivity readings were taken. A set of control samples were kept on ice at 4°C for the duration of the freezing portion of the experiment. Percent electrolyte leakage (%EL) was calculated for each sample as the conductivity prior to being autoclaved (i.e., cell damage due to freezing) divided by the conductivity after autoclaving (i.e., total cell lysis) multiplied by 100. As the leakage of electrolytes in the control samples is not caused by freezing, we normalized the % EL of each sample (% $EL_{sample}$) to the average % EL of the control samples (% $EL_{control}$) and to a maximum electrolyte leakage of 100 %. For this purpose, we used the % EL of the lowest freezing temperature (% $EL_{max}$). We normalized electrolyte leakage using the equation:

$$\%EL_{Norm} = \frac{\%EL_{Sample} - mean(\%EL_{Control})}{\%EL_{Max} - mean(\%EL_{Control})} \times 100$$

EL values of samples contaminated by liquid in the waterbaths were removed to avoid biased results. Non-linear least squares logistic models were fit to the gametophyte and sporophyte data independently and compared to a model with both sets of data and the Bayesian Information Criterion (BIC) for the models were compared.

http://www.repeatmasker.org

26. Kim D, Paggi JM, Park C, Bennett C, Salzberg SL. Graph-based genome alignment and genotyping with HISAT2 and HISAT-genotype. Nat Biotechnol. 2019 Aug 2;37(8):907–15.
27. Merchant SS, Prochnik SE, Vallon O, Harris EH, Karpowicz SJ, Witman GB, et al. The Chlamydomonas genome reveals the evolution of key animal and plant functions. Science. 2007 Oct 12;318(5848):245–50.
28. Li F-W, Nishiyama T, Waller M, Frangedakis E, Keller J, Li Z, et al. *Anthoceros* genomes illuminate the origin of land plants and the unique biology of hornworts. Nat Plants. 2020 Mar 13;6(3):259–72.
29. Bowman JL, Kohchi T, Yamato KT, Jenkins J, Shu S, Ishizaki K, et al. Insights into Land Plant Evolution Garnered from the *Marchantia polymorpha* Genome. Cell. 2017 Oct 5;171(2):287-304.e15.
30. Rensing SA, Lang D, Zimmer AD, Terry A, Salamov A, Shapiro H, et al. The Physcomitrella genome reveals evolutionary insights into the conquest of land by plants. Science. 2008 Jan 4;319(5859):64–9.
31. Banks JA, Nishiyama T, Hasebe M, Bowman JL, Gribskov M, dePamphilis C, et al. The *Selaginella* genome identifies genetic changes associated with the evolution of vascular plants. Science. 2011 May 20;332(6032):960–3.
32. Li F-W, Brouwer P, Carretero-Paulet L, Cheng S, de Vries J, Delaux P-M, et al. Fern genomes elucidate land plant evolution and cyanobacterial symbioses. Nat Plants. 2018 Jul 2;4(7):460–72.
33. Rahmatpour N, Kuo L-Y, Kang J, Herman E, Lei L, Li M, et al. Analyses of *Marsilea vestita* genome and transcriptomes do not support widespread intron retention during spermatogenesis. New Phytol. 2023 Mar;237(5):1490–4.
34. Marchant DB, Chen G, Cai S, Chen F, Schafran P, Jenkins J, et al. Dynamic genome evolution in a model fern. Nat Plants. 2022 Sep 1;8(9):1038–51.
35. Fang Y, Qin X, Liao Q, Du R, Luo X, Zhou Q, et al. The genome of homosporous maidenhair fern sheds light on the euphyllophyte evolution and defences. Nat Plants. 2022 Sep 1;8(9):1024–37.
36. Huang X, Wang W, Gong T, Wickell D, Kuo L-Y, Zhang X, et al. The flying spider-monkey tree fern genome provides insights into fern evolution and arborescence. Nat Plants. 2022 May 9;8(5):500–12.
37. Amborella Genome Project. The Amborella genome and the evolution of flowering plants. Science. 2013 Dec 20;342(6165):1241089.
38. Kawahara Y, de la Bastide M, Hamilton JP, Kanamori H, McCombie WR, Ouyang S, et al. Improvement of the *Oryza sativa* Nipponbare reference genome using next generation sequence and optical map data. Rice (N Y). 2013 Feb 6;6(1):4.
39. Hufford MB, Seetharam AS, Woodhouse MR, Chougule KM, Ou S, Liu J, et al. De novo assembly, annotation, and comparative analysis of 26 diverse maize genomes. Science. 2021 Aug 6;373(6555):655–62.
40. Tuskan GA, Difazio S, Jansson S, Bohlmann J, Grigoriev I, Hellsten U, et al. The genome of black cottonwood, *Populus trichocarpa* (Torr. & Gray). Science. 2006 Sep 15;313(5793):1596–604.
41. Arabidopsis Genome Initiative. Analysis of the genome sequence of the flowering plant *Arabidopsis thaliana*. Nature. 2000 Dec 14;408(6814):796–815.
42. Gabriel L, Brůna T, Hoff KJ, Ebel M, Lomsadze A, Borodovsky M, et al. BRAKER3: Fully automated genome annotation using RNA-seq and protein evidence with GeneMark-ETP, AUGUSTUS, and TSEBRA. Genome Res. 2024 Jun 25;34(5):769–77.
43. Stanke M, Schöffmann O, Morgenstern B, Waack S. Gene prediction in eukaryotes with a generalized hidden Markov model that uses hints from external sources. BMC Bioinformatics. 2006 Feb 9;7:62.
44. Stanke M, Diekhans M, Baertsch R, Haussler D. Using native and syntenically mapped cDNA alignments to improve *de novo* gene finding. Bioinformatics. 2008 Mar 1;24(5):637–44.
45. Buchfink B, Xie C, Huson DH. Fast and sensitive protein alignment using DIAMOND. Nat Methods. 2015 Jan;12(1):59–60.
46. Brůna T, Lomsadze A, Borodovsky M. GeneMark-ETP significantly improves the accuracy of automatic annotation of large eukaryotic genomes. Genome Res. 2024 Jun 25;34(5):757–68.
47. Pertea G, Pertea M. GFF utilities: gffread and gffcompare. F1000Res. 2020 Apr 28;9:304.
48. Gotoh O. A space-efficient and accurate method for mapping and aligning cDNA sequences onto genomic

sequence. Nucleic Acids Res. 2008 May;36(8):2630–8.
49. Iwata H, Gotoh O. Benchmarking spliced alignment programs including Spaln2, an extended version of Spaln that incorporates additional species-specific features. Nucleic Acids Res. 2012 Nov 1;40(20):e161.
50. Kovaka S, Zimin AV, Pertea GM, Razaghi R, Salzberg SL, Pertea M. Transcriptome assembly from long-read RNA-seq alignments with StringTie2. Genome Biol. 2019 Dec 16;20(1):278.
51. Nevers Y, Warwick Vesztrocy A, Rossier V, Train C-M, Altenhoff A, Dessimoz C, et al. Quality assessment of gene repertoire annotations with OMArk. Nat Biotechnol. 2024 Feb 21;
52. Dainat J, Hereñú D. NBISweden/AGAT: AGAT-v0.4.0. Zenodo. 2020;
53. Jones P, Binns D, Chang H-Y, Fraser M, Li W, McAnulla C, et al. InterProScan 5: genome-scale protein function classification. Bioinformatics. 2014 May 1;30(9):1236–40.
54. Cantalapiedra CP, Hernández-Plaza A, Letunic I, Bork P, Huerta-Cepas J. eggNOG-mapper v2: Functional annotation, orthology assignments, and domain prediction at the metagenomic scale. Mol Biol Evol. 2021 Dec 9;38(12):5825–9.
55. Huerta-Cepas J, Szklarczyk D, Heller D, Hernández-Plaza A, Forslund SK, Cook H, et al. eggNOG 5.0: a hierarchical, functionally and phylogenetically annotated orthology resource based on 5090 organisms and 2502 viruses. Nucleic Acids Res. 2019 Jan 8;47(D1):D309–14.
56. Bolger AM, Lohse M, Usadel B. Trimmomatic: A flexible trimmer for Illumina sequence data. Bioinformatics. 2014 Aug 1;30(15):2114–20.
57. McKenna A, Hanna M, Banks E, Sivachenko A, Cibulskis K, Kernytsky A, et al. The Genome Analysis Toolkit: a MapReduce framework for analyzing next-generation DNA sequencing data. Genome Res. 2010 Sep;20(9):1297–303.
58. Akalin A, Kormaksson M, Li S, Garrett-Bakelman FE, Figueroa ME, Melnick A, et al. methylKit: a comprehensive R package for the analysis of genome-wide DNA methylation profiles. Genome Biol. 2012 Oct 3;13(10):R87.
59. R Core Team. R: A language and environment for statistical computing. Vienna, Austria: R Foundation for Statistical Computing; 2024.
60. Huang X, Zhang S, Li K, Thimmapuram J, Xie S, Wren J. ViewBS: a powerful toolkit for visualization of high-throughput bisulfite sequencing data. Bioinformatics. 2018 Feb 15;34(4):708–9.
61. Takuno S, Gaut BS. Body-methylated genes in Arabidopsis thaliana are functionally important and evolve slowly. Mol Biol Evol. 2012 Jan;29(1):219–27.
62. Lister R, O'Malley RC, Tonti-Filippini J, Gregory BD, Berry CC, Millar AH, et al. Highly integrated single-base resolution maps of the epigenome in Arabidopsis. Cell. 2008 May 2;133(3):523–36.
63. Takuno S, Gaut BS. Gene body methylation is conserved between plant orthologs and is of evolutionary consequence. Proc Natl Acad Sci USA. 2013 Jan 29;110(5):1797–802.
64. Niederhuth CE, Bewick AJ, Ji L, Alabady MS, Kim KD, Li Q, et al. Widespread natural variation of DNA methylation within angiosperms. Genome Biol. 2016 Sep 27;17(1):194.
65. Fay MP, Shaw PA. Exact and Asymptotic Weighted Logrank Tests for Interval Censored Data: The interval R package. J Stat Softw. 2010 Aug;36(2).
66. Alexa A, Rahnenfuhrer J. topGO: Enrichment Analysis for Gene Ontology. BioConductor; 2020.
67. Robinson MD, McCarthy DJ, Smyth GK. edgeR: a Bioconductor package for differential expression analysis of digital gene expression data. Bioinformatics. 2010 Jan 1;26(1):139–40.
68. Li C, Wickell D, Kuo L-Y, Chen X, Nie B, Liao X, et al. Extraordinary preservation of gene collinearity over three hundred million years revealed in homosporous lycophytes. Proc Natl Acad Sci USA. 2024 Jan 23;121(4):e2312607121.
69. Wickell D, Kuo L-Y, Yang H-P, Dhabalia Ashok A, Irisarri I, Dadras A, et al. Underwater CAM photosynthesis elucidated by *Isoetes* genome. Nat Commun. 2021 Nov 3;12(1):6348.
70. Breinholt JW, Carey SB, Tiley GP, Davis EC, Endara L, McDaniel SF, et al. A target enrichment probe set for resolving the flagellate land plant tree of life. Appl Plant Sci. 2021 Jan 24;9(1):e11406.
71. Camacho C, Coulouris G, Avagyan V, Ma N, Papadopoulos J, Bealer K, et al. BLAST+: architecture and

applications. BMC Bioinformatics. 2009 Dec 15;10:421.
72. Ranwez V, Harispe S, Delsuc F, Douzery EJP. MACSE: Multiple Alignment of Coding SEquences accounting for frameshifts and stop codons. PLoS ONE. 2011 Sep 16;6(9):e22594.
73. Capella-Gutiérrez S, Silla-Martínez JM, Gabaldón T. trimAl: A tool for automated alignment trimming in large-scale phylogenetic analyses. Bioinformatics. 2009 Aug 1;25(15):1972–3.
74. Kalyaanamoorthy S, Minh BQ, Wong TKF, von Haeseler A, Jermiin LS. ModelFinder: fast model selection for accurate phylogenetic estimates. Nat Methods. 2017 Jun;14(6):587–9.
75. Minh BQ, Schmidt HA, Chernomor O, Schrempf D, Woodhams MD, von Haeseler A, et al. IQ-TREE 2: New models and efficient methods for phylogenetic inference in the genomic era. Mol Biol Evol. 2020 May 1;37(5):1530–4.
76. Hoang DT, Chernomor O, von Haeseler A, Minh BQ, Vinh LS. Ufboot2: improving the ultrafast bootstrap approximation. Mol Biol Evol. 2018 Feb 1;35(2):518–22.
77. Zhang C, Rabiee M, Sayyari E, Mirarab S. ASTRAL-III: Polynomial time species tree reconstruction from partially resolved gene trees. BMC Bioinformatics. 2018 May 8;19(Suppl 6):153.
78. Smith SA, O'Meara BC. treePL: Divergence time estimation using penalized likelihood for large phylogenies. Bioinformatics. 2012 Oct 15;28(20):2689–90.
79. Morris JL, Puttick MN, Clark JW, Edwards D, Kenrick P, Pressel S, et al. The timescale of early land plant evolution. Proc Natl Acad Sci USA. 2018 Mar 6;115(10):E2274–83.
80. Putri GH, Anders S, Pyl PT, Pimanda JE, Zanini F. Analysing high-throughput sequencing data in Python with HTSeq 2.0. Bioinformatics. 2022 Mar 21;
81. Love MI, Huber W, Anders S. Moderated estimation of fold change and dispersion for RNA-seq data with DESeq2. Genome Biol. 2014;15(12):550.
82. Ogata H, Goto S, Sato K, Fujibuchi W, Bono H, Kanehisa M. KEGG: kyoto encyclopedia of genes and genomes. Nucleic Acids Res. 1999 Jan 1;27(1):29–34.
83. Wu T, Hu E, Xu S, Chen M, Guo P, Dai Z, et al. clusterProfiler 4.0: A universal enrichment tool for interpreting omics data. Innovation (Camb). 2021 Aug 28;2(3):100141.
84. Emms DM, Kelly S. OrthoFinder: solving fundamental biases in whole genome comparisons dramatically improves orthogroup inference accuracy. Genome Biol. 2015 Aug 6;16(1):157.
85. Emms DM, Kelly S. OrthoFinder: phylogenetic orthology inference for comparative genomics. Genome Biol. 2019 Nov 14;20(1):238.
86. Berardini TZ, Reiser L, Li D, Mezheritsky Y, Muller R, Strait E, et al. The *Arabidopsis* information resource: making and mining the "gold standard" annotated reference plant genome. Genesis. 2015 Aug 4;53(8):474–85.
87. Mukherjee K, Brocchieri L, Bürglin TR. A comprehensive classification and evolutionary analysis of plant homeobox genes. Mol Biol Evol. 2009 Dec;26(12):2775–94.
88. Rozewicki J, Li S, Amada KM, Standley DM, Katoh K. MAFFT-DASH: integrated protein sequence and structural alignment. Nucleic Acids Res. 2019 Jul 2;47(W1):W5–10.
89. Chamala S, Feng G, Chavarro C, Barbazuk WB. Genome-wide identification of evolutionarily conserved alternative splicing events in flowering plants. Front Bioeng Biotechnol. 2015 Mar 26;3:33.
90. Mei W, Boatwright L, Feng G, Schnable JC, Barbazuk WB. Evolutionarily conserved alternative splicing across monocots. Genetics. 2017 Oct;207(2):465–80.
91. Li H, Handsaker B, Wysoker A, Fennell T, Ruan J, Homer N, et al. The Sequence Alignment/Map format and SAMtools. Bioinformatics. 2009 Aug 15;25(16):2078–9.
92. Danecek P, Bonfield JK, Liddle J, Marshall J, Ohan V, Pollard MO, et al. Twelve years of SAMtools and BCFtools. Gigascience. 2021 Feb 16;10(2).
93. Pertea M, Pertea GM, Antonescu CM, Chang T-C, Mendell JT, Salzberg SL. StringTie enables improved reconstruction of a transcriptome from RNA-seq reads. Nat Biotechnol. 2015 Mar;33(3):290–5.
94. Hass BJ. Analysis of Alternative Splicing in Plants with Bioinformatics Tools. In: Reddy ASN, Golovkin M, editors. Nuclear pre-mRNAProcessing i n Plants. Springer; 2008. p. 17–37.
95. Li H. Minimap2: pairwise alignment for nucleotide sequences. Bioinformatics. 2018 Sep 15;34(18):3094–100.

96. Philips SJ, Dudík M, Schapire RE. Maxent software for modeling species niches and distributions. American Musuem of Natural History; 2024.
97. Hijmans RJ, Philips S, Leathwick J, Elith J. dismo. CRAN; 2024.
98. Fick SE, Hijmans RJ. WorldClim 2: new 1-km spatial resolution climate surfaces for global land areas. Int J Climatol. 2017 Oct;37(12):4302–15.
99. Suissa JS, Sundue MA, Testo WL. Mountains, climate and niche heterogeneity explain global patterns of fern diversity. J Biogeogr. 2021 Feb 16;
100. Tatebe H, Ogura T, Nitta T, Komuro Y, Ogochi K, Takemura T, et al. Description and basic evaluation of simulated mean state, internal variability, and climate sensitivity in MIROC6. Geosci Model Dev. 2019 Jul 8;12(7):2727–65.
101. Thalhammer A, Hincha DK, Zuther E. Measuring freezing tolerance: electrolyte leakage and chlorophyll fluorescence assays. Methods Mol Biol. 2014;1166:15–24.
102. Kovaleski AP, Grossman JJ. Standardization of electrolyte leakage data and a novel liquid nitrogen control improve measurements of cold hardiness in woody tissue. Plant Methods. 2021 May 22;17(1):53.

**SUPPLEMENTAL FIGURES**

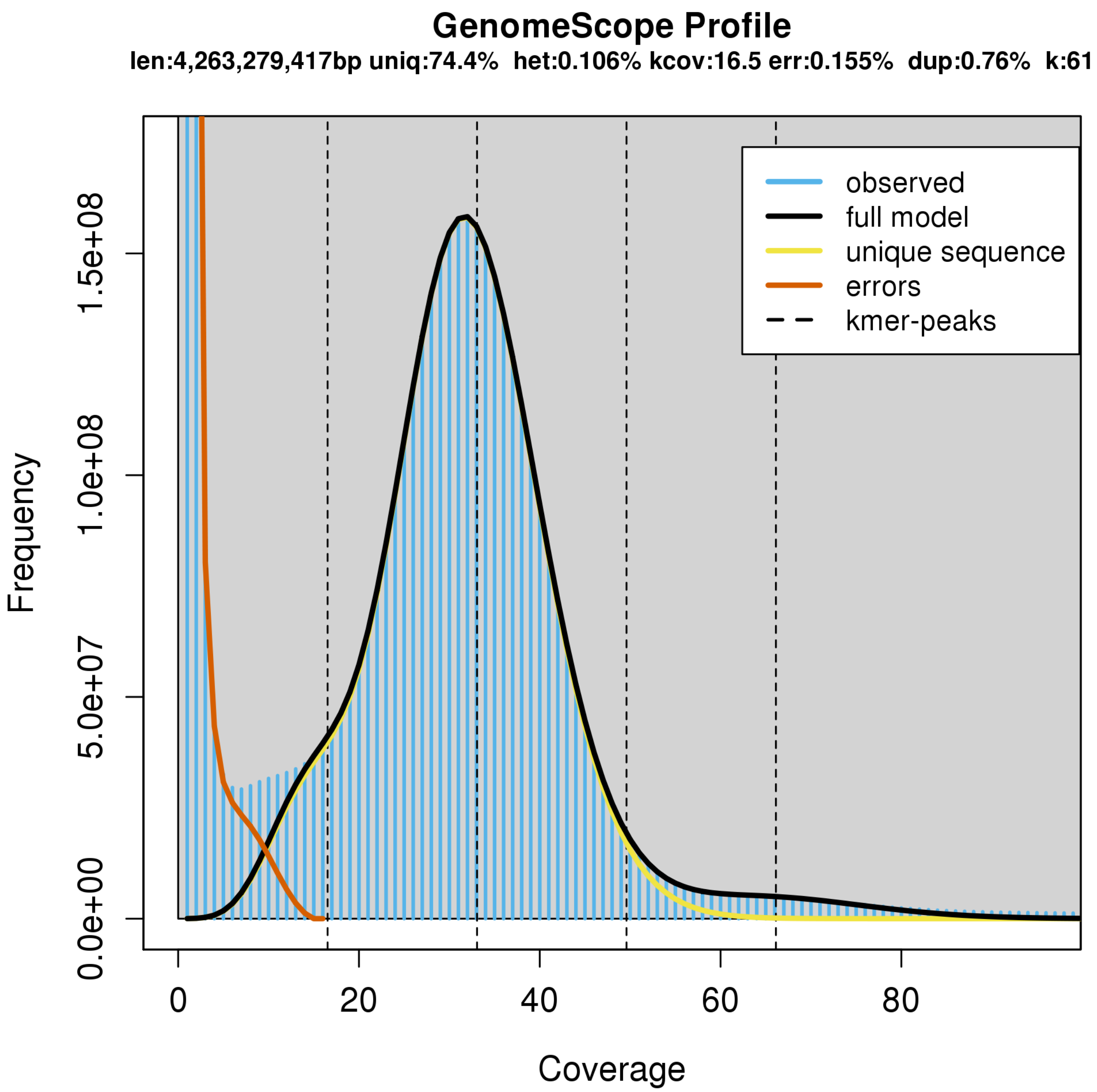


**Figure S1.** Genomescope results with 61-mers from short-read Illumina data.

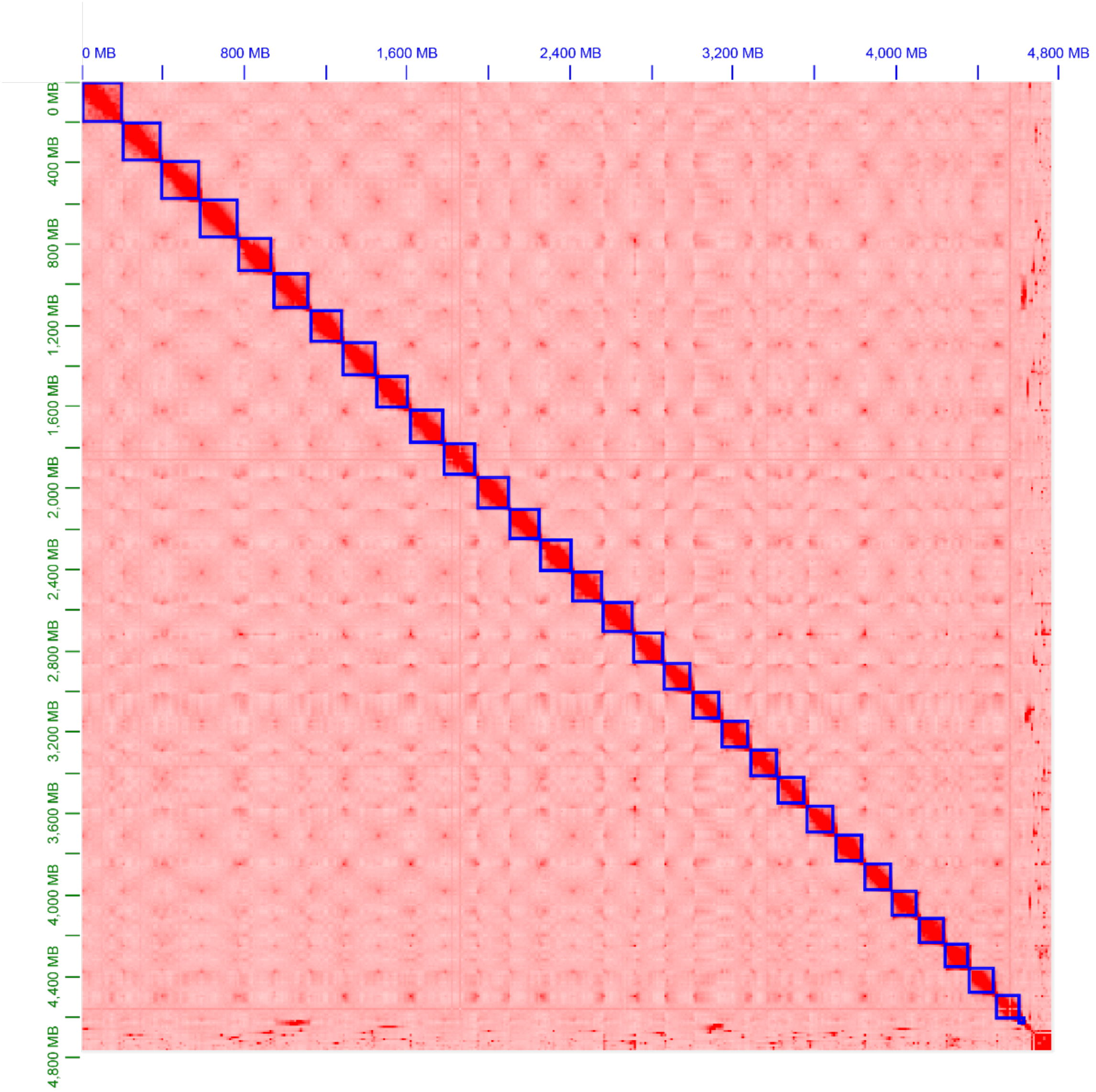


**Figure S2.** Hi-C contact map showing 30 chromosome-level scaffolds.

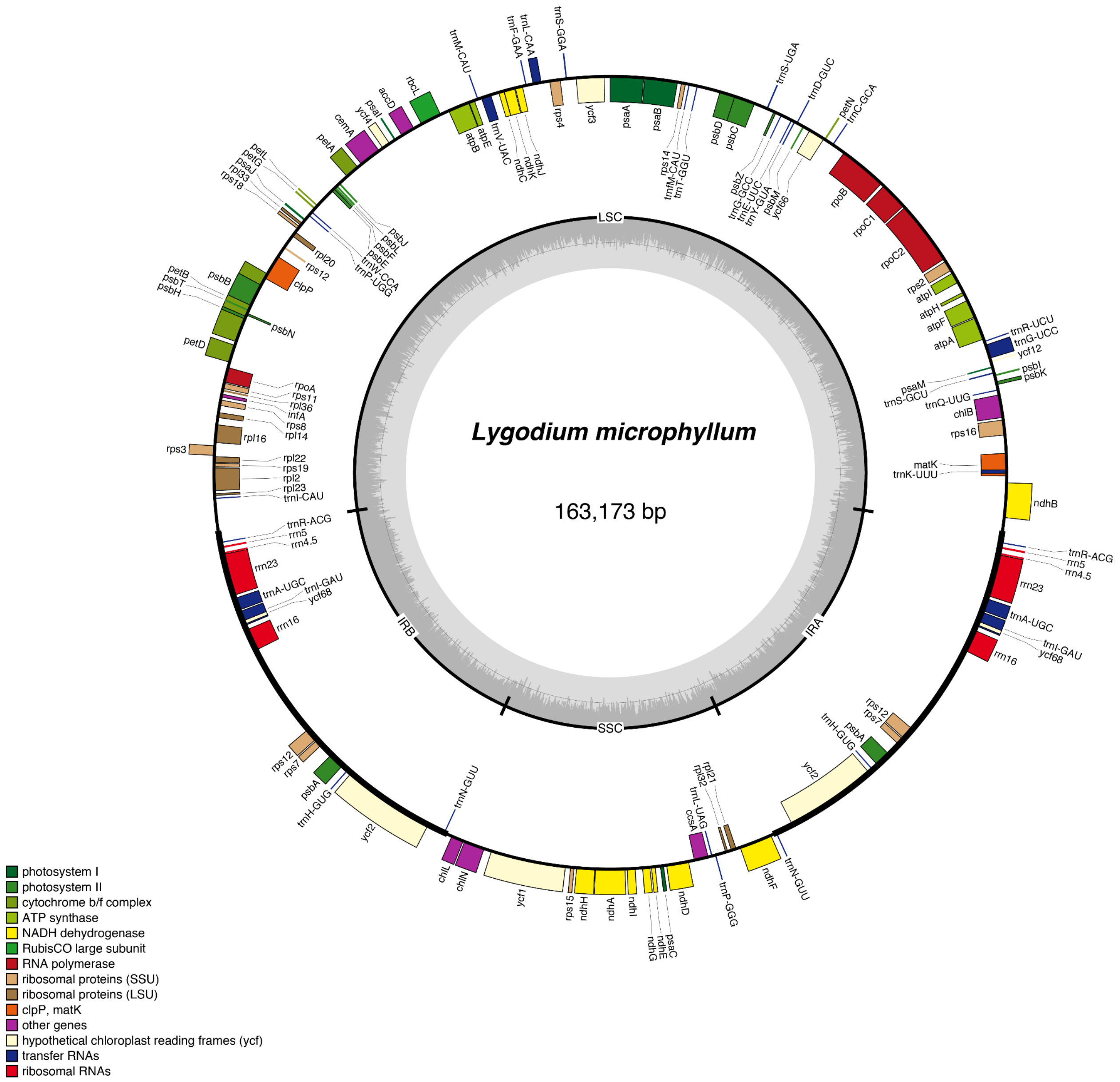


**Figure S3.** *Lygodium microphyllum* plastome assembly and annotation.

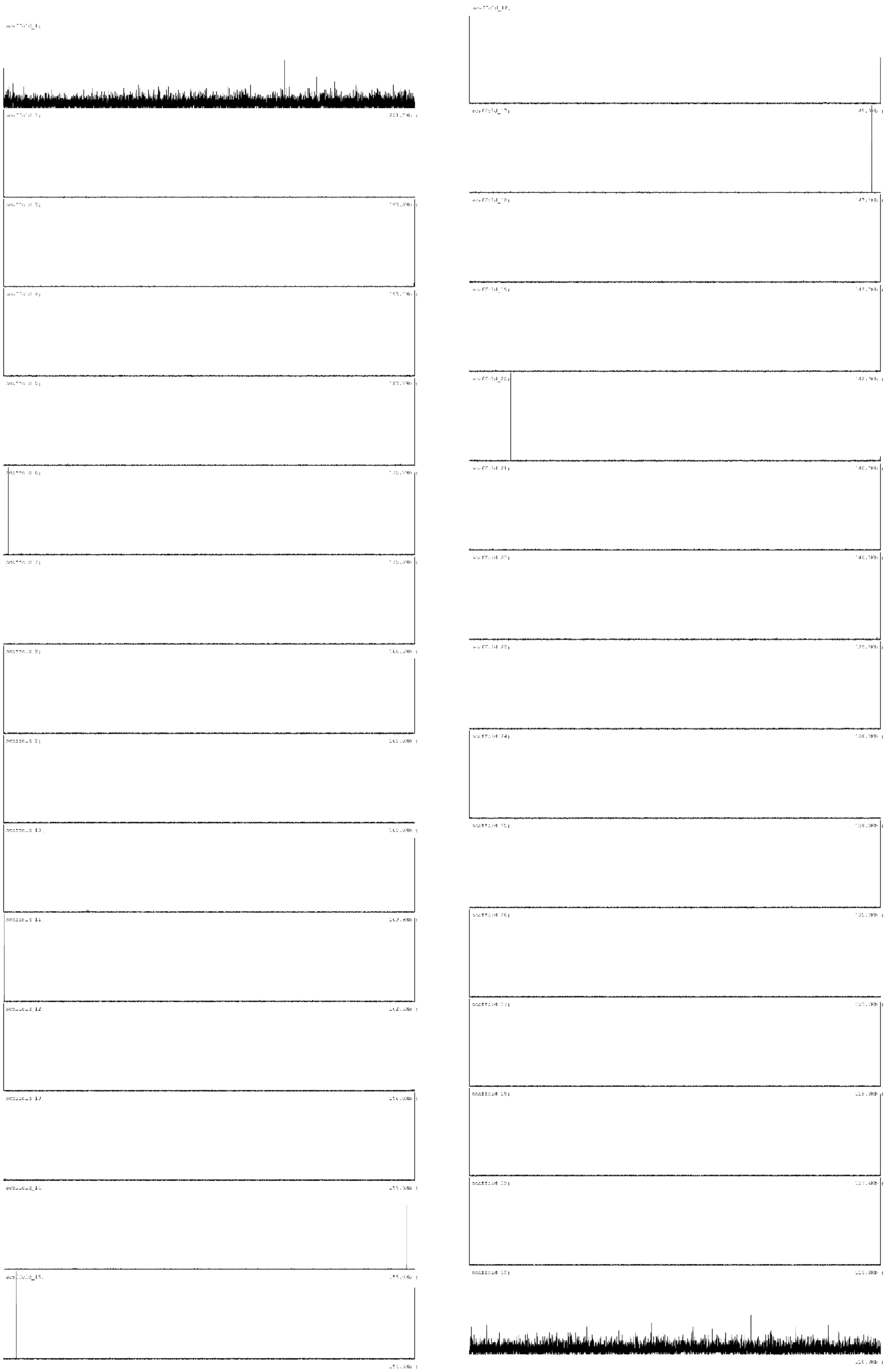

**Figure S4.** Telomeric repeat frequency across 30 chromosome-level scaffolds.

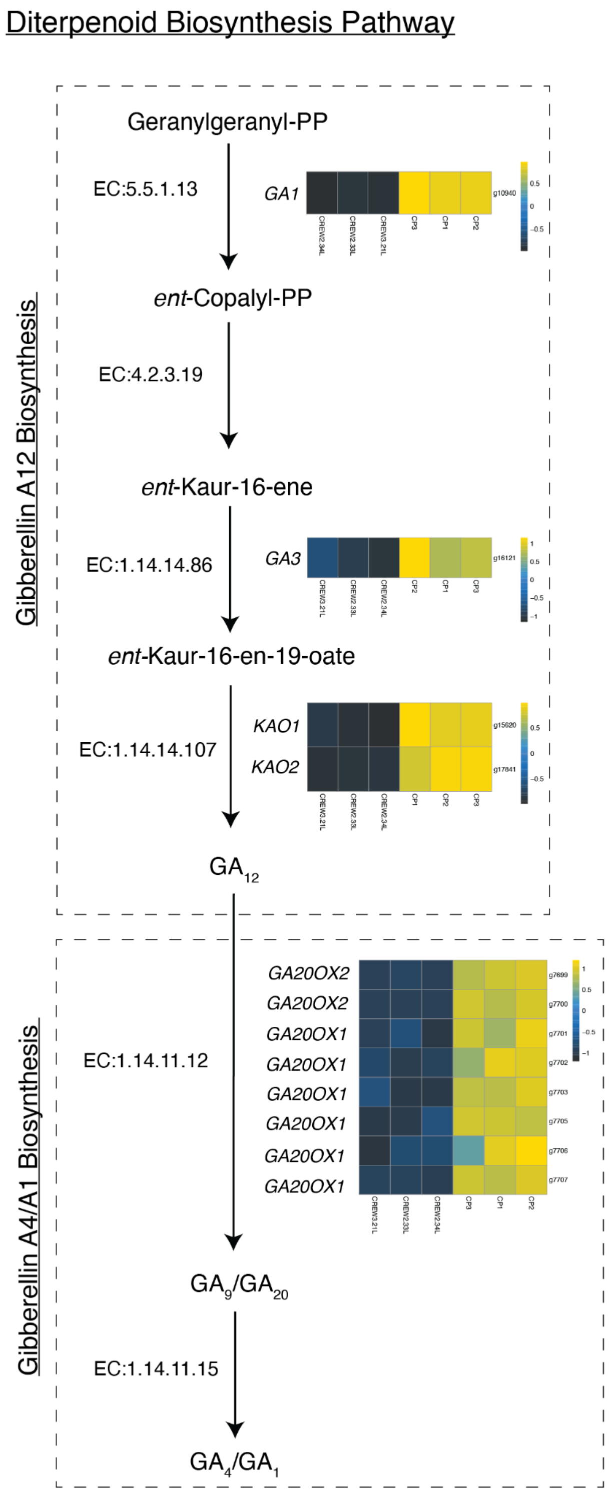


**Figure S5.** Selected portion of enriched KEGG pathway "ath00904" diterpenoid biosynthesis in upregulated genes in the gametophyte relative to the leaf. The gibberellin A12 biosynthesis and gibberellin A4/A1 biosynthesis modules were enriched within the pathway. Heatmaps show Z-scores of significantly differentially expressed genes involved in reactions. Pathways without heatmaps did not have significantly differentially expressed genes and are therefore not shown.

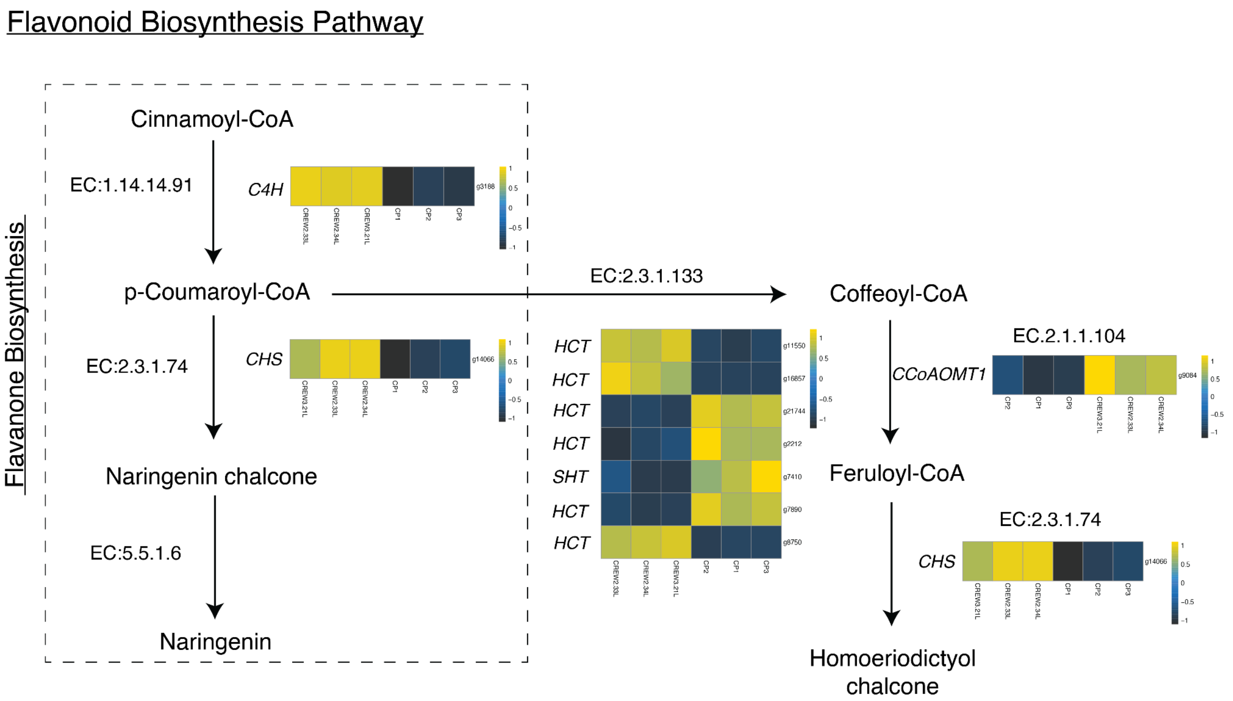


**Figure S6.** Selected portion of enriched KEGG pathway "ath00941" flavonoid biosynthesis in downregulated genes in the gametophyte relative to the leaf. The flavanone biosynthesis module was enriched within the pathway. Heatmaps show Z-scores of significantly differentially expressed genes involved in reactions. Pathways without heatmaps did not have significantly differentially expressed genes and are therefore not shown.

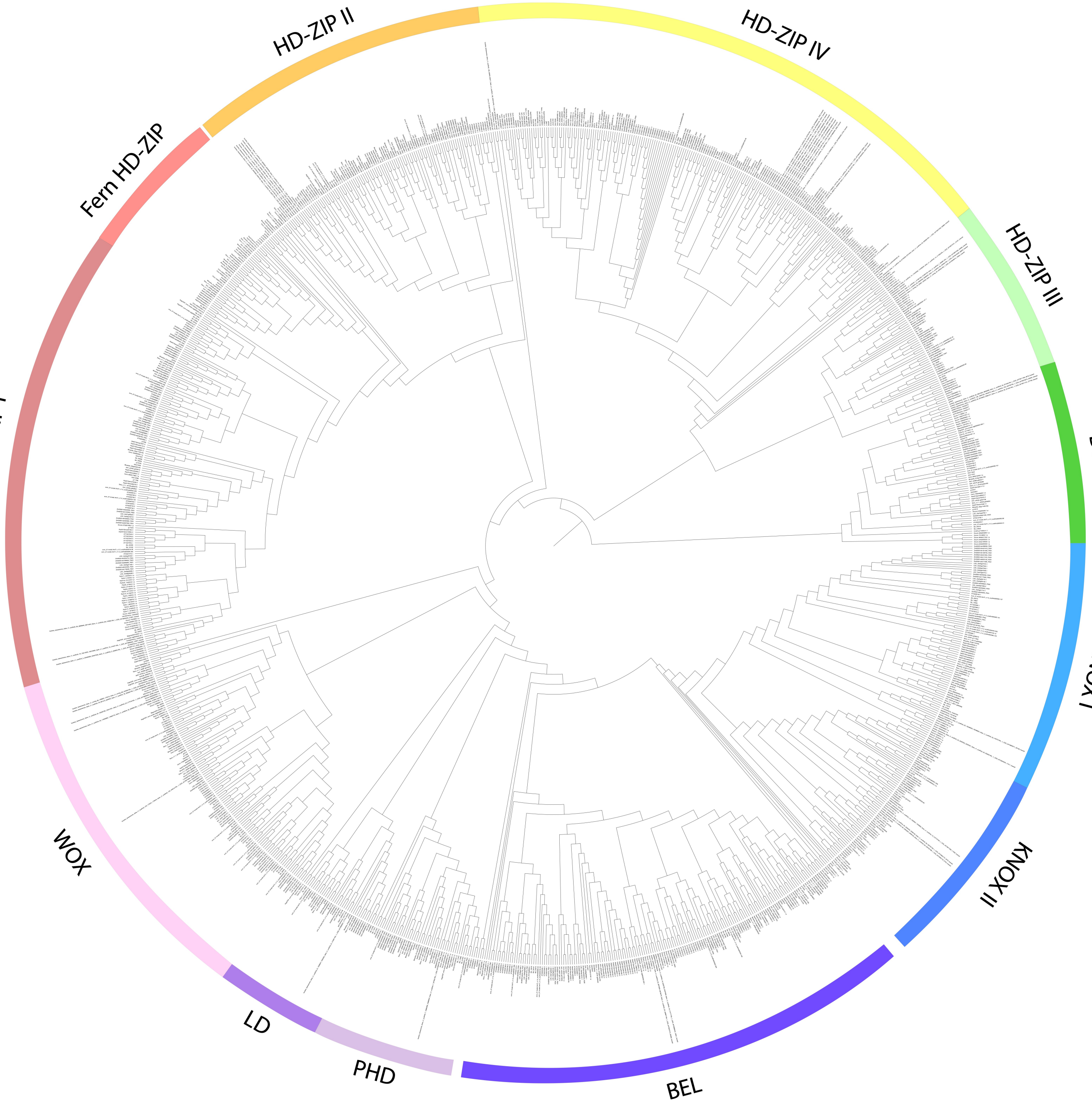


**Figure S7.** Phylogeny of homeobox genes across land plants.

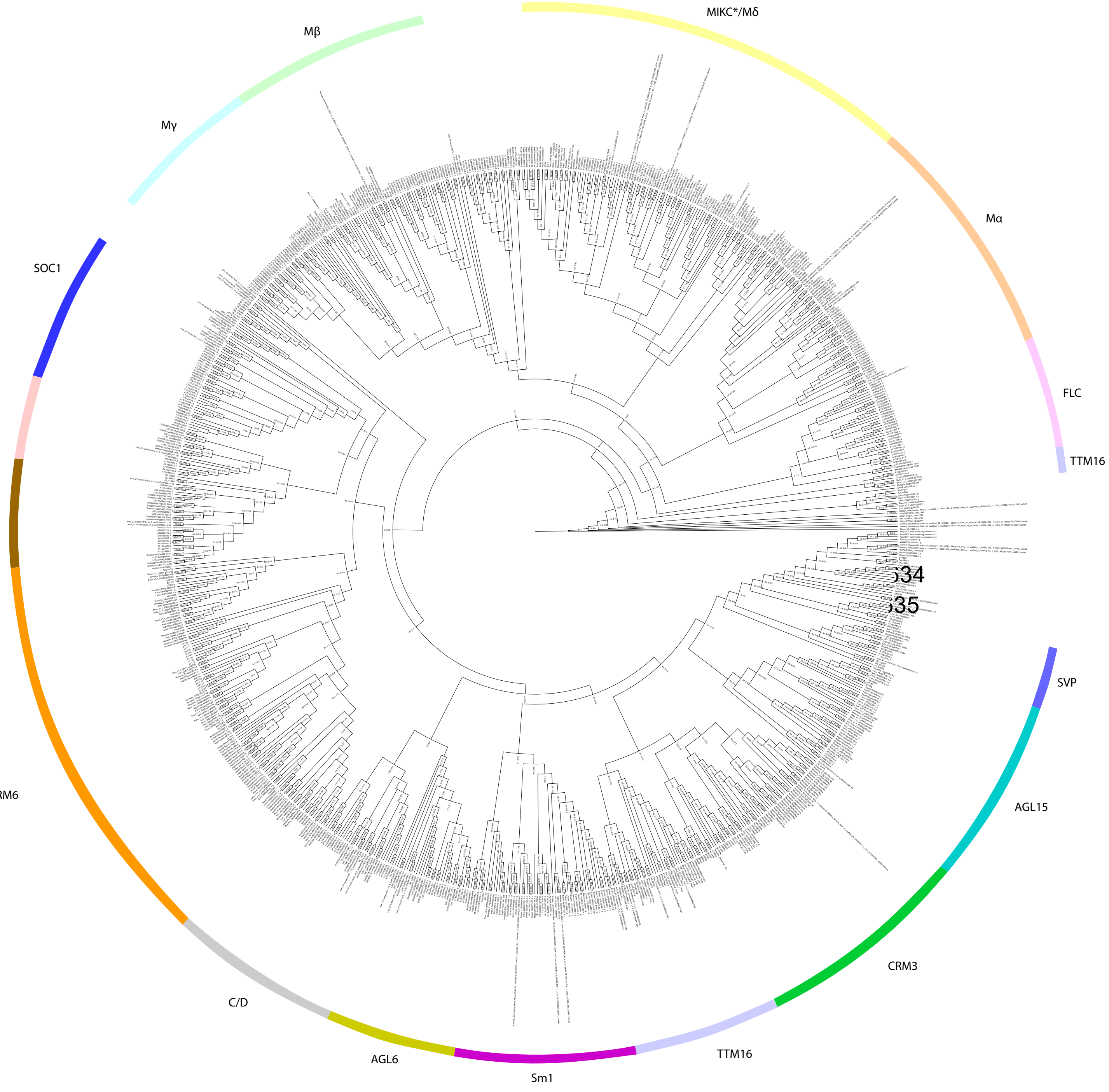


**Figure S8.** Phylogeny of MADS-box genes across land plants. Node values are ultra-fast bootstrap values / SHaLRT values.

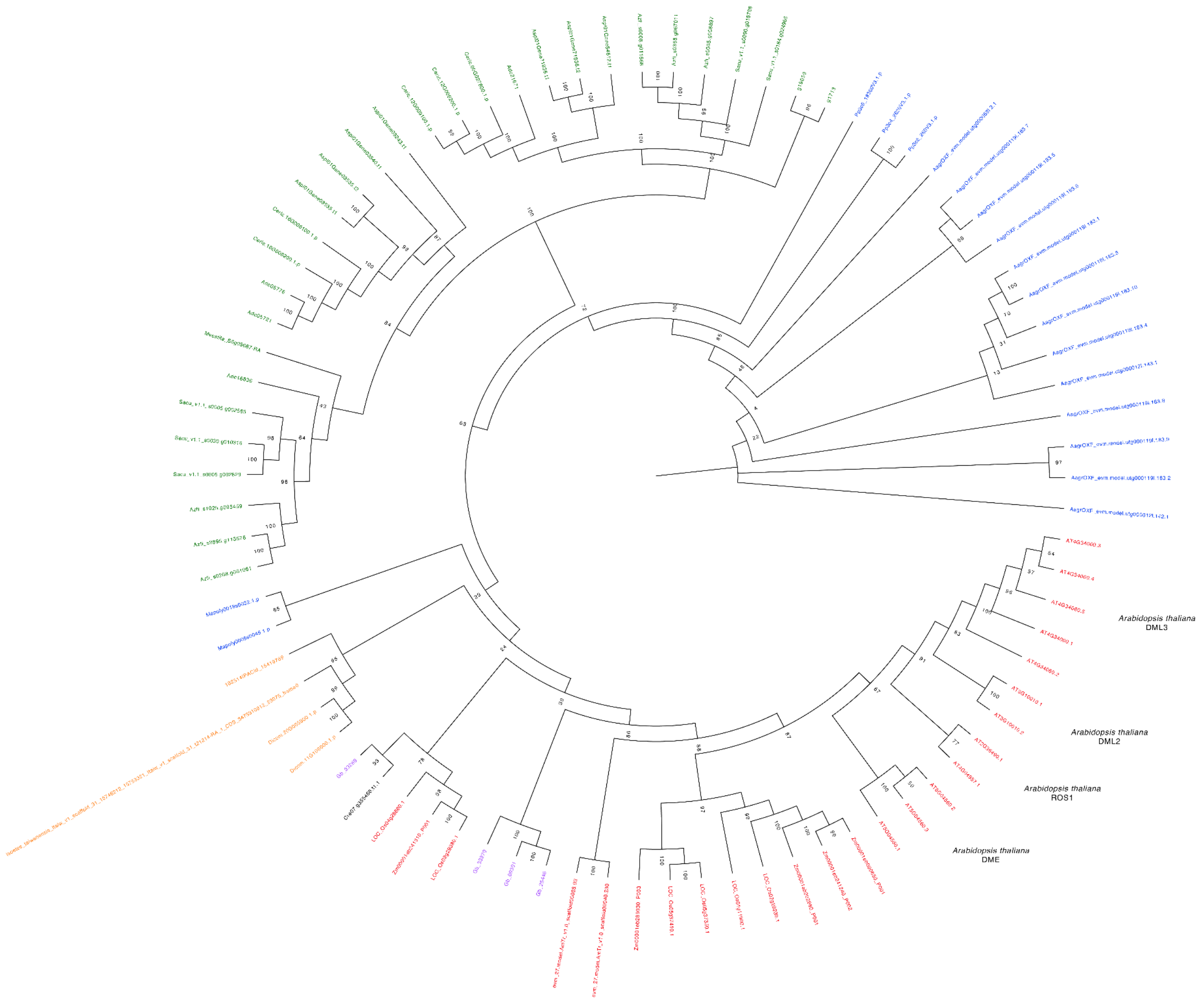


**Figure S9.** Phylogeny of DNA demethylation genes (DME, ROS1, DML2, DML3). Node values are bootstrap metrics from 1000 ultrafast bootstraps. Tips are colored based on lineage: hornworts are blue, lycophytes are orange, ferns are green, gymnosperms are purple, angiosperms are red.

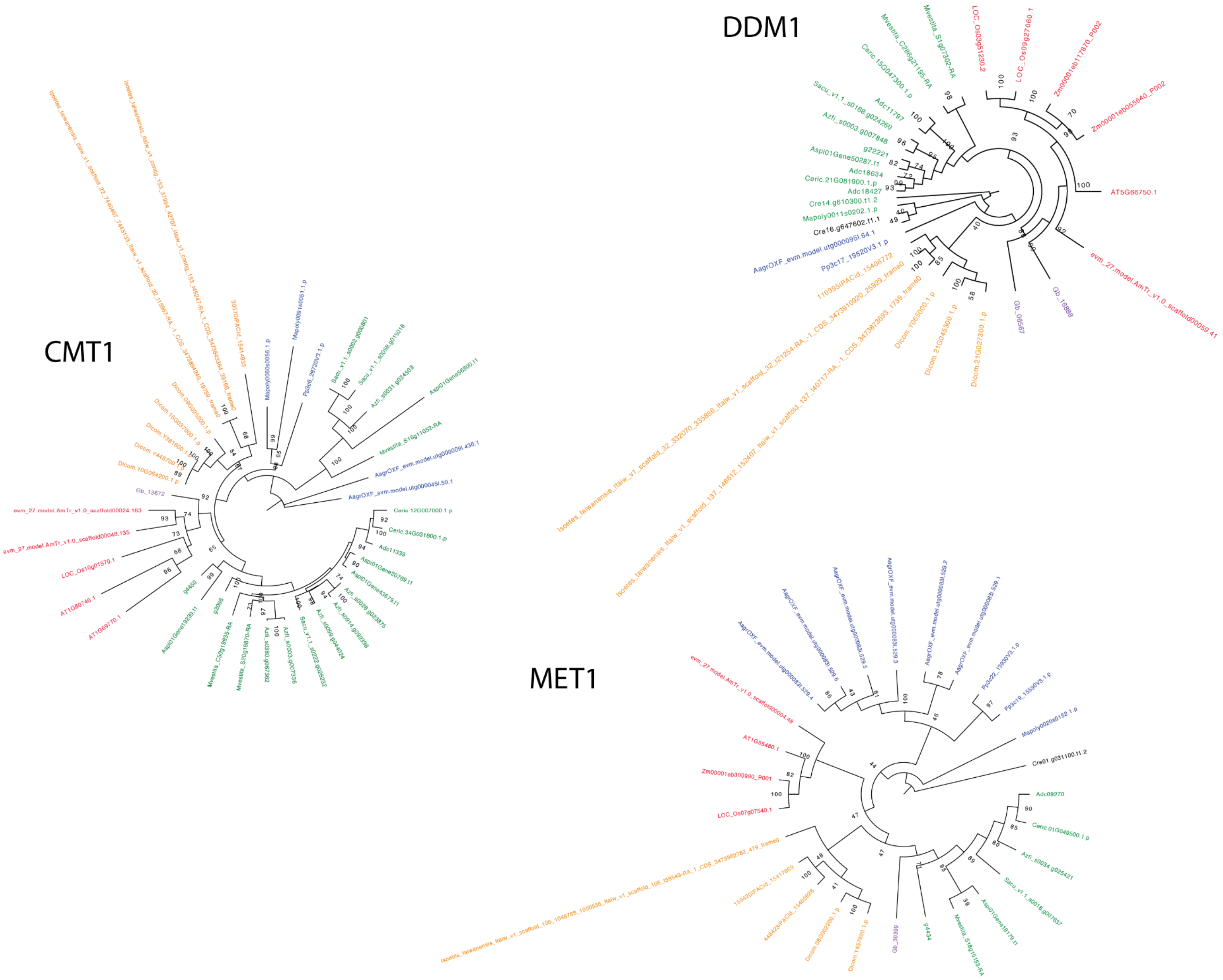


**Figure S10.** Phylogeny of genes involved in DNA methylation maintenance (CMT1, DDM1, MET1 . Node values are bootstrap metrics from 1000 ultrafast bootstraps. Tips are colored based on lineage: hornworts are blue, lycophytes are orange, ferns are green, gymnosperms are purple, angiosperms are red.

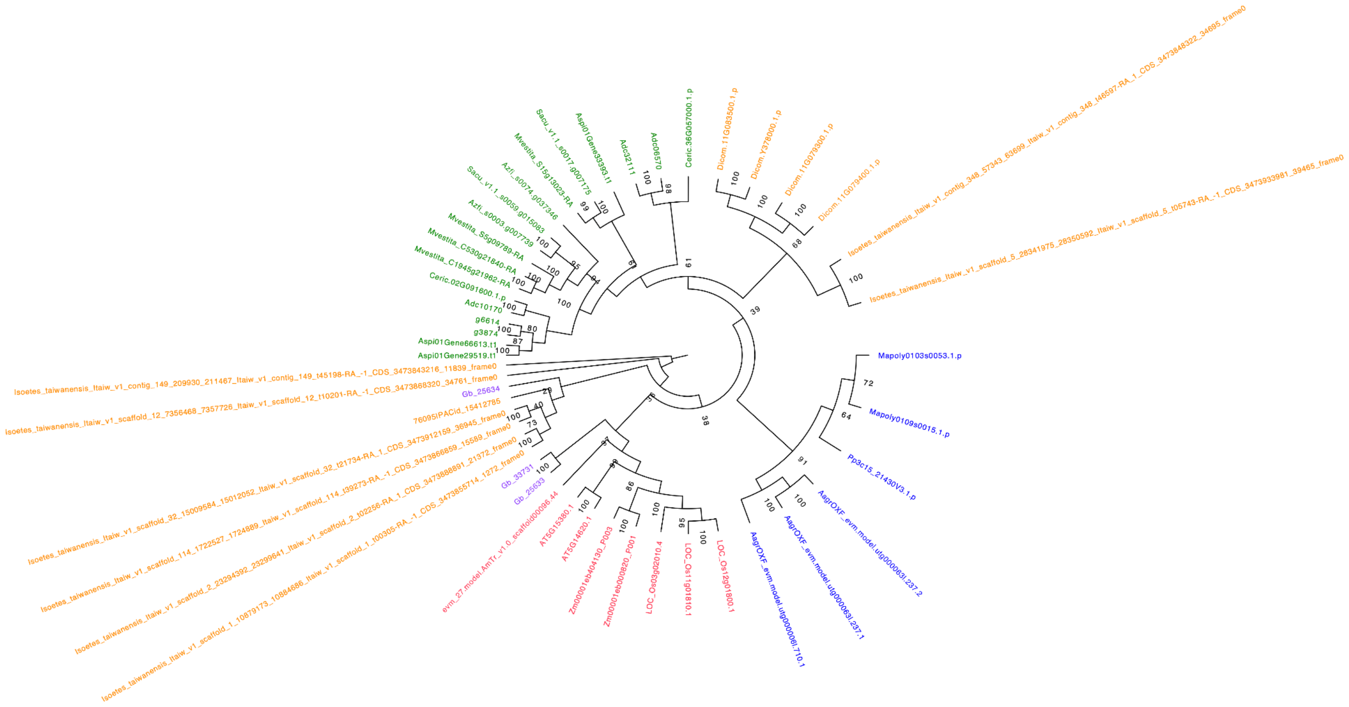

**Figure S11.** Phylogeny of *de novo* DNA methylation genes (DRM1, DRM2). Node values are bootstrap metrics from 1000 ultrafast bootstraps. Tips are colored based on lineage: hornworts are blue, lycophytes are orange, ferns are green, gymnosperms are purple, angiosperms are red.

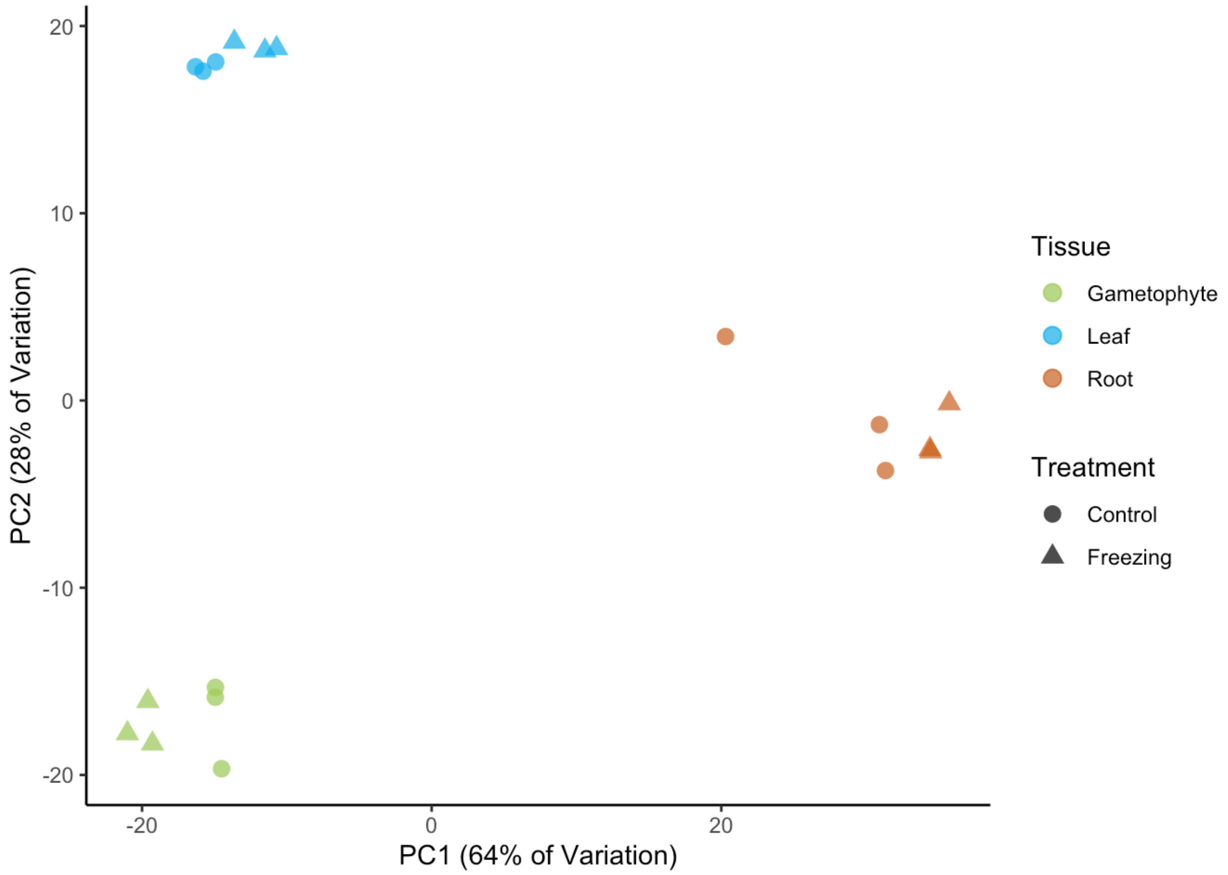


**Figure S12.** PCA of significant differentially expressed genes following variance stabilizing transformation for each tissue and treatment type in the freezing experiment.

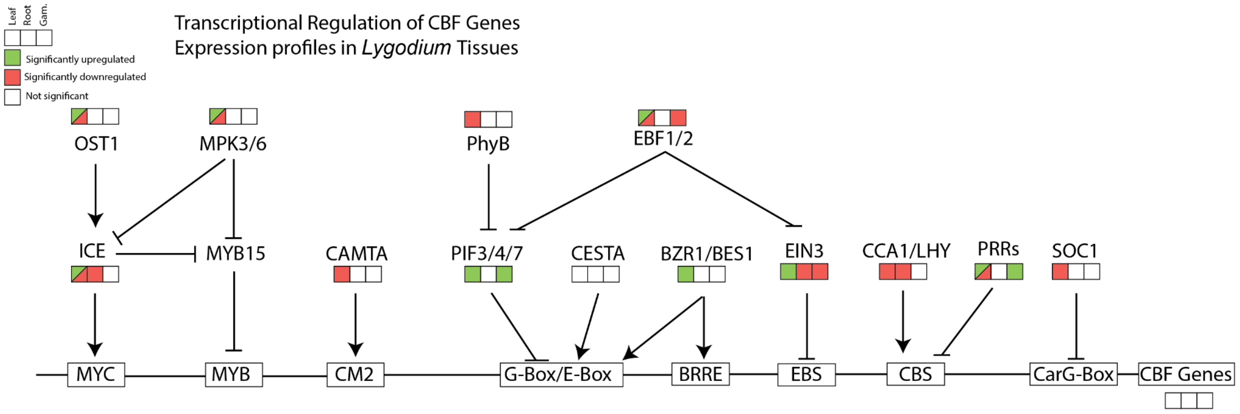


**Figure S13.** Transcriptional regulation of CBF genes in *Lygodium* root, gametophyte, and leaf tissues as a response to freezing shock. From left to right, boxes show significant upregulation (green) or downregulation (red) in the leaf, root, and gametophyte tissues.

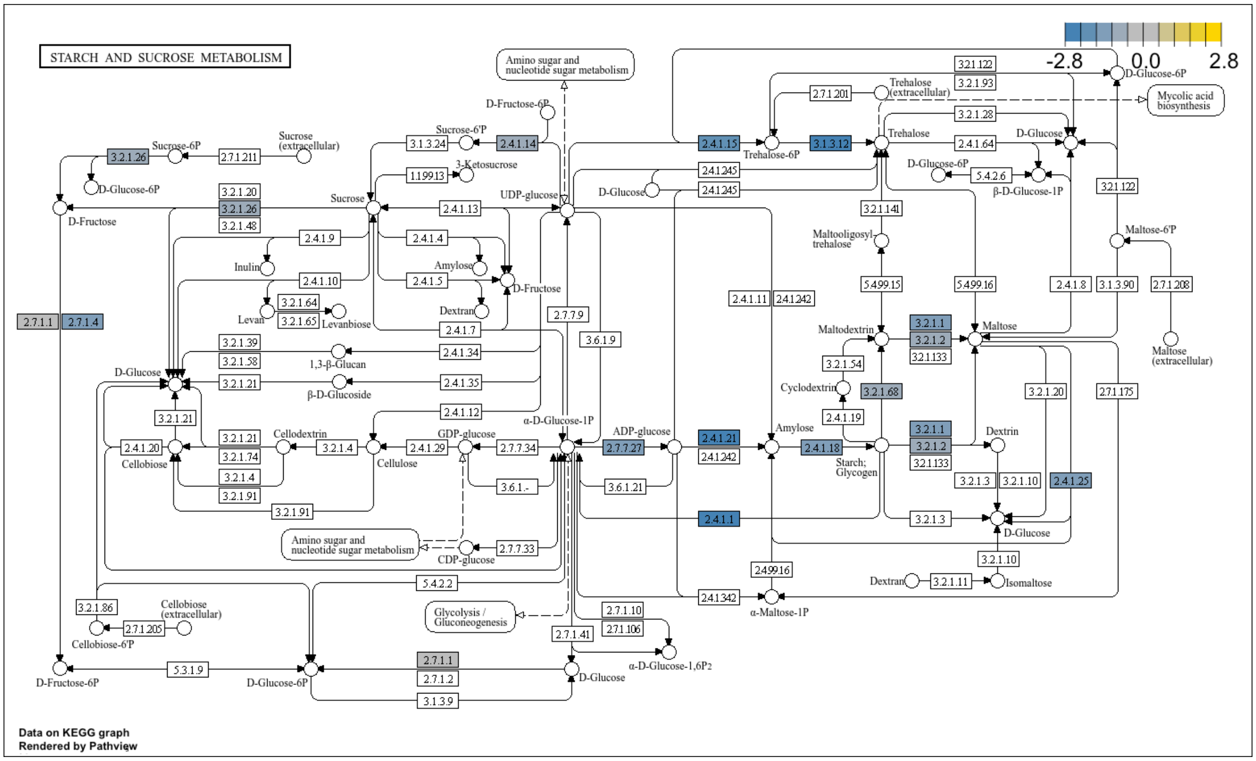

**Figure S14.** Enriched KEGG pathway "ath00500" in downregulated genes in the leaf due to freezing. Colored boxes show specific enriched enzymatic reactions, colored by log2foldchange in expression.

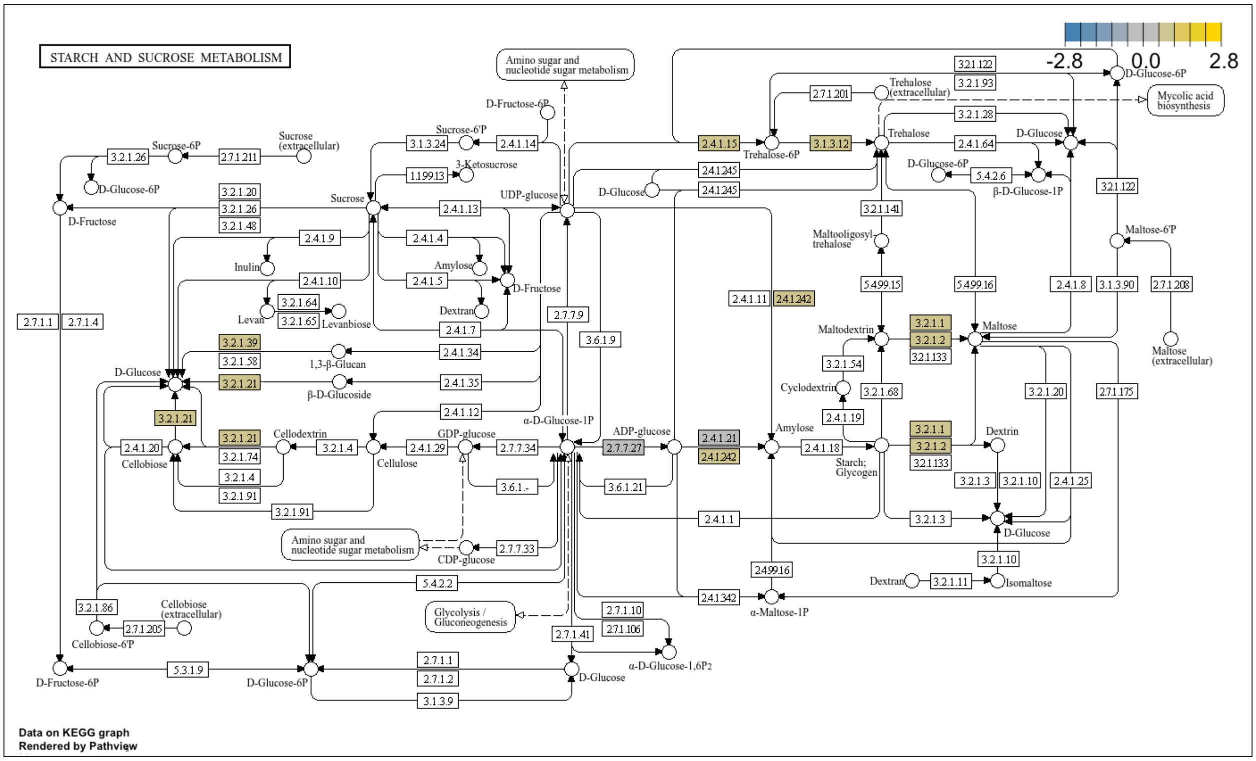


**Figure S15.** Enriched KEGG pathway "ath00500" in upregulated genes in the gametophyte due to freezing. Colored boxes show specific enriched enzymatic reactions, colored by log2foldchange in expression.

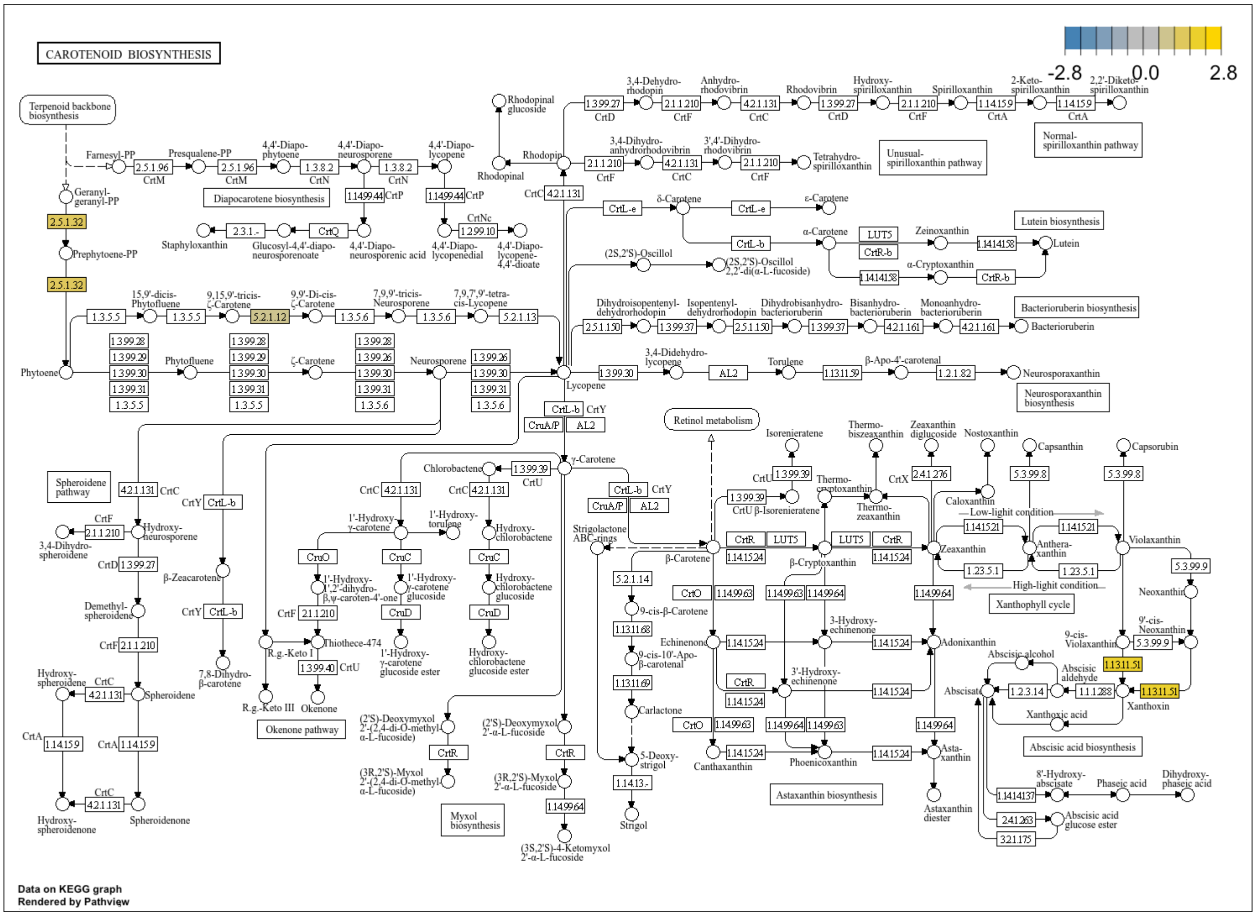


**Figure S16.** Enriched KEGG pathway "ath00906" in upregulated genes in the root due to freezing. Colored boxes show specific enriched enzymatic reactions, colored by log2foldchange in expression.

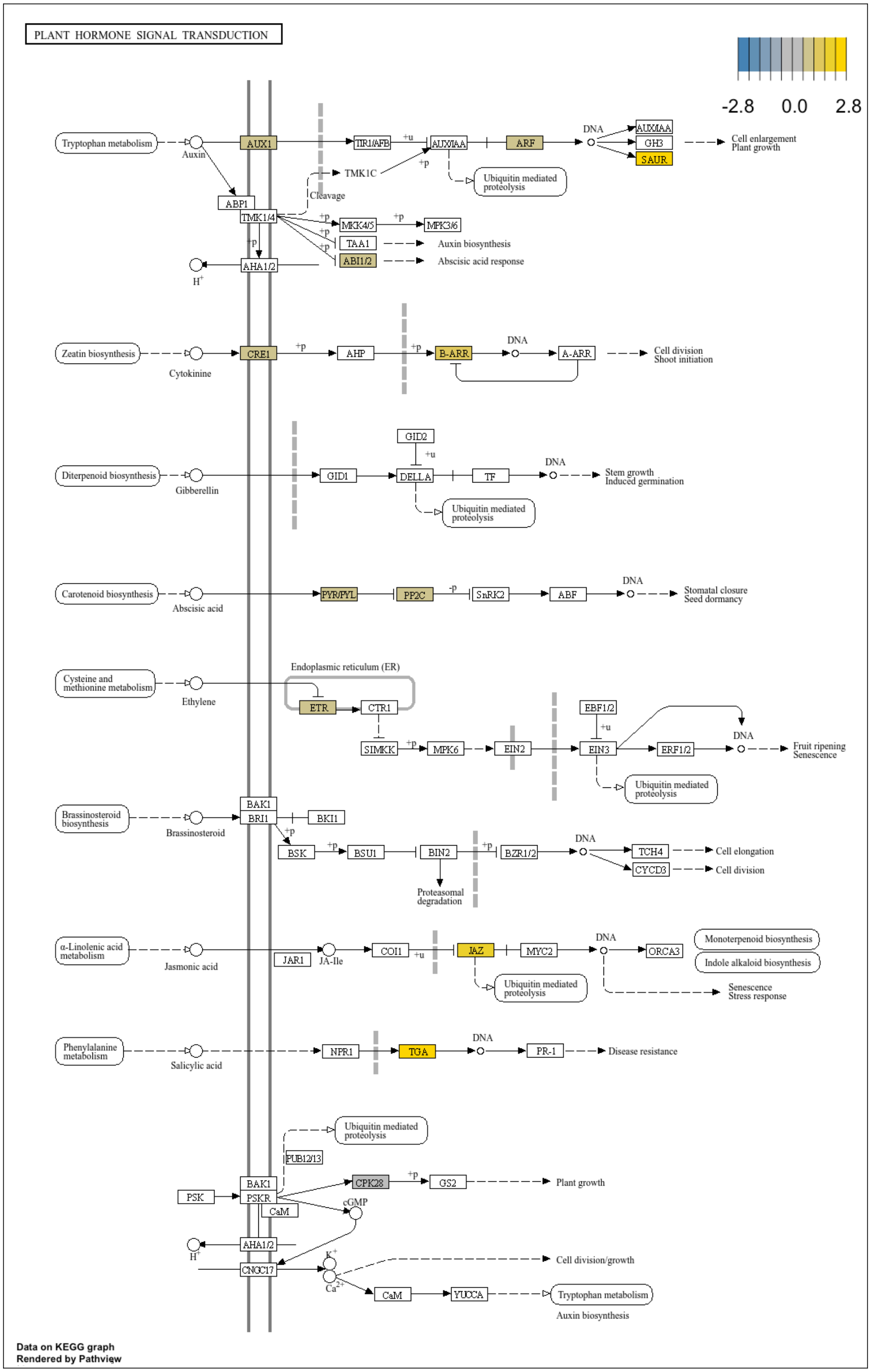


**Figure S17.** Enriched KEGG pathway "ath04705" in upregulated genes in the root due to freezing. Colored boxes show specific enriched enzymatic reactions, colored by log2foldchange in expression.
